# Spliceosomal complex components are critical for adjusting the C:N balance during high-light acclimation

**DOI:** 10.1101/2023.07.19.549727

**Authors:** Gali Estopare Araguirang, Benedikt Venn, Nadja-Magdalena Kelber, Regina Feil, John Lunn, Tatjana Kleine, Dario Leister, Timo Mühlhaus, Andreas S. Richter

## Abstract

Plant acclimation to an ever-changing environment is decisive for growth, reproduction and survival. Light availability limits biomass production on both ends of the intensity spectrum. Therefore, the adjustment of plant metabolism is central to high-light (HL) acclimation, and accumulation of photoprotective anthocyanins is commonly observed. However, mechanisms and factors regulating the HL acclimation response are less clear. Two Arabidopsis mutants of spliceosome components exhibiting a pronounced anthocyanin overaccumulation in HL were isolated from a forward genetic screen for new factors crucial for plant acclimation. Time-resolved physiological, transcriptome and metabolome analysis revealed a vital function of the spliceosome components for rapidly adjusting gene expression and metabolism. Deficiency of INCREASED LEVEL OF POLYPLOIDY1 (ILP1), NTC-RELATED PROTEIN1 (NTR1), and PLEIOTROPIC REGULATORY LOCUS1 (PRL1) resulted in a marked overaccumulation of carbohydrates and strongly diminished amino acid biosynthesis in HL. While not generally limited in N-assimilation, *ilp1*, *ntr1,* and *prl1* mutants showed higher glutamate levels and reduced amino acid biosynthesis in HL. The comprehensive analysis reveals a function of the spliceosome in the conditional regulation of the carbon:nitrogen-balance and the accumulation of anthocyanins during HL acclimation. The importance of gene expression, metabolic regulation, and re-direction of carbon towards anthocyanin biosynthesis for HL acclimation are discussed.

## Introduction

Plants are remarkable organisms capable of acclimating to changes in their environment. Among others, changes in light intensity and growth temperature are factors potentially deleterious for plant growth, reproduction and survival. Plants employ various mechanisms to thrive under diverse light conditions, including high light (HL) acclimation. HL acclimation allows plants to adjust their physiological and biochemical responses to cope with excessive light exposure (Bailey et al., 2002; Garcia-Molina et al., 2020; Kleine et al., 2021). When exposed to intense light, plants face the challenge of excessive energy absorption, leading to the activation of alternative energy dissipation and electron transfer routes within the photosynthetic complexes, eventually resulting in photoinhibition and production of harmful reactive oxygen species (ROS) and subsequent damage to cellular components (Mittler et al., 2022). The HL acclimation response of plants allows the dissipation of excess energy, protects photosynthetic machinery, maintains cellular homeostasis, and permits metabolic reprogramming (Joliot and Johnson, 2011; Goss and Lepetit, 2015; Huang et al., 2019; Balcke et al., 2023). One notable aspect of HL acclimation in plants is the biosynthesis of anthocyanins. Beyond their aesthetic appeal, anthocyanins serve critical biological and eco-physiological functions, such as attracting pollinators, protecting against UV radiation, and acting as antioxidants (Gould, 2004; Gould et al., 2018; Agati et al., 2020; Agati et al., 2021).

Anthocyanin biosynthesis is a complex and highly regulated process that involves multiple enzymatic reactions across different cellular compartments (as reviewed by Araguirang and Richter, 2022). After the conversion of chloroplast-derived phenylalanine to *p*-coumaroyl CoA via reactions of phenylpropanoid biosynthesis, early biosynthetic genes, such as chalcone synthase, provide the basic flavan skeleton to different branches of flavonoid biosynthesis. Enzymes involved in anthocyanin biosynthesis are collectively summarised as late biosynthetic genes acting on different dihydro flavonols (Vogt, 2010; Saito et al., 2013). Dihydroflavonol 4-reductase (DFR) yields leucocyanidins which serve as substrates for anthocyanin biosynthesis through methyl-, acyl and glycosyl transferases. The expressions of early and late biosynthesis genes is under tight control of different transcription factors (TFs). The early biosynthesis genes are regulated by *MYB11*, *MYB12,* and *MYB111* (Stracke et al., 2007). On the other hand, late biosynthesis genes, essential for anthocyanin accumulation, are controlled by an MBW complex [<u>M</u>YB, <u>b</u>asic helix-loop-helix (bHLH), <u>W</u>D40]. The WD40 variant, *TRANSPARENT TESTA GLABRA 1 (TTG1*), together with bHLH TFs, *TRANSPARENT TESTA 8* (*TT8*), *GLABRA 3* (*GL3*), and *ENHANCER OF GL3* (*EGL3)*, act in concert with MYB TFs such as *MYB75,* also known as *PRODUCTION OF ANTHOCYANIN PIGMENT 1* (*PAP1*), *MYB90* (*PAP2*), *MYB113*, and *MYB114* (Borevitz et al., 2000; Gonzalez et al., 2008; Lloyd et al., 2017). Over the years, several factors have been proposed that initiate the production of anthocyanins in HL, many of which involve the chloroplasts as a source for a retrograde signal regulating the expression of anthocyanin biosynthesis genes (Araguirang and Richter, 2022; Richter et al., 2023). For instance, the *genomes uncoupled* (*gun*) chloroplast-to-nucleus communication pathway is essential to coordinate the expression of photosynthesis and anthocyanin biosynthesis genes during chloroplast biogenesis. A mutant for the Mg-chelatase H subunit vital for chlorophyll biosynthesis, *gun5-1,* showed perturbed induction of anthocyanin biosynthesis when chloroplast biogenesis was blocked by norflurazon (Richter et al., 2020). Besides a potential influence of ROS and phytohormones, we previously found that photosynthetic activity, connected carbon fixation and increased cellular sugar contents in HL are the primary signal to induce anthocyanin biosynthesis in HL. Furthermore, the activation of anthocyanin biosynthesis requires inactivation of the SNF1-RELATED KINASE 1 (SnRK1), most likely through regulatory sugar phosphates such as trehalose 6-phosphate (Tre6P) (Zirngibl et al., 2023). In addition to anthocyanin biosynthesis, biosynthesis of nitrogen (N) containing compounds, such as amino acids and proteins, is stimulated when increased photosynthetic activity provides more carbon skeletons to plant metabolism in HL (Garcia-Molina et al., 2020; Balcke et al., 2023). Hence, metabolic acclimation maintains the C:N balance and properly distributes available resources to different biochemical pathways. While metabolic acclimation requires fast regulation of enzymes on the post-translational level (for example, redox-regulation, Dietz and Hell, 2015), the plant acclimation response to environmental factors is largely also underpinned by the adjustment of gene expression (Huang et al., 2019; Garcia-Molina et al., 2020).

One essential component for the maturation of primary transcripts is a highly dynamic supramolecular ribonucleoprotein complex and intricate machinery known as the spliceosome. The spliceosome, which is highly conserved among eukaryotes, plays a fundamental role in pre-mRNA splicing, a crucial step in gene expression that enables the precise removal of non-coding introns and splicing together of coding exons (Black, 2003; Wahl et al., 2009). The assembled spliceosome has at least seven consecutive states during the whole splicing event, namely the pre-catalytic, activated, step I catalytically activated, a catalytic step I, step II catalytically activated, post-catalytic, and the intron lariat spliceosome (ILS) (Wan et al., 2017). The ILS is disassembled during the last stage of the canonical splicing cycle and thus releasing splicing factors to be recycled and the intron lariats to be degraded (Arenas and Abelson, 1997). Deficiencies of two critical components of the ILS disassembly complex, *INCREASED LEVEL OF POLYPLOIDY 1* (*ILP1*) and *NTC-RELATED PROTEIN 1*/*SPLICEOSOMAL TIMEKEEPER LOCUS 1* (*NTR1*/*STIPL1*) have been shown to cause perturbation of the circadian clock, cell cycle, and heat stress tolerance, primarily because of alternative splicing (Yoshizumi et al., 2006; Jones et al., 2012; Dolata et al., 2015; Wang et al., 2019; Cecchini et al., 2022; He et al., 2023).

Alternative splicing is a critical process in eukaryotes that allows a single gene to produce multiple protein isoforms by selectively joining different exons during mRNA processing (Kan et al., 2001; Brett et al., 2002). This process greatly expands the proteome diversity and can significantly affect protein accumulation and regulation (Syed et al., 2012). It can influence various aspects of protein function, including enzymatic activity, stability, localisation, and interaction partners (Lamberto et al., 2010; Kashkan et al., 2022). The precise regulation of alternative splicing is also crucial for cellular functions and developmental processes and dependent on environmental cues like light signals, cold stress, and high temperature (Fu et al., 2009; Calixto et al., 2018; John et al., 2021; Martín et al., 2021; Martín, 2023). That is why in NTR1- and ILP1-deficient mutants, hundreds of genes are differentially spliced, and the plants consequently display pleiotropic phenotypes, including a reduction in microRNA (miRNA) levels (Dolata et al., 2015; Wang et al., 2019).

MiRNAs are small non-coding RNA molecules that play crucial roles in post-transcriptional gene silencing (Lee and Ambros, 2001; Ruvkun, 2001). They bind to specific target mRNA molecules, leading to degradation or translational repression. The intricate relationship between splicing and miRNA accumulation involves the direct and physical interaction of ILP1 and NTR1 with SERRATE, a core protein in miRNA biogenesis, in the nucleus (Wang et al., 2019). Previous studies have shown that alternative splicing and miRNAs regulate anthocyanin biosynthesis (Li et al., 2019; Chen et al., 2020; Wang et al., 2020). However, how the ILS disassembly factors and, to an extent, the whole spliceosomal complex adjust and maintain the metabolic balance during HL acclimation remains largely unknown.

Here we show that the spliceosomal components ILP1, and Pleiotropic Regulatory Locus 1 (PRL1), isolated from a forward genetic screen for factors regulating anthocyanin biosynthesis, and NTR1 play regulatory roles in the expression and processing of mRNAs amino acid, carbohydrate and anthocyanin biosynthesis and maintenance of the C:N balance during HL acclimation. We propose that the involvement of the spliceosomal complex in metabolic regulation is essential to provide a new level of regulation of the HL acclimation response.

## Results

### Isolation and characterisation of *ilp1* mutant by forward genetic screening

A forward genetic screen was conceived to isolate new factors regulating anthocyanin biosynthesis in response to plastid-derived factors (Fig. S1). The suppressor screen was based on the *gun5-1* mutant perturbed in retrograde signalling when plastid biogenesis is suppressed, resulting in a pronounced anthocyanin deficiency when germinated on media supplemented with Norflurazon (NF) (Richter et al., 2020). NF inhibits carotenoid biosynthesis, resulting in white-to-purplish plants devoid of photosynthetic pigments and visible anthocyanin pigmentation (Fig. S1A). After ethyl methanesulfonate mutagenesis, M2 seeds were germinated in the presence of norflurazon and screened for *gun5-1* mutants showing a *restored anthocyanin accumulation* (*raa*) phenotype. Several *raa* mutants were obtained, and the *raa20, raa7 and raa14* mutants were selected for detailed characterisation (Fig. S1A). We found that perturbation of *RAA20* in the background of *gun5-1* also resulted in a more pronounced accumulation of anthocyanins compared to the corresponding wild-type (WT) accession (Col-0) and the parental *gun5-1* mutant after 24 h HL exposure (Fig. S1A), suggesting that RAA20 function is also involved in the HL acclimation response of plants. Whole genome sequencing of *raa20* individuals re-selected in the T2 after backcrossing to *gun5-1* revealed n SNP at position 2767004 of chromosome 5, which is the first nucleotide of intron 4, causing retention of intron 4 in the *ILP1* mRNA (AT5G08550) (Fig. S1B-D, Supplemental Table S7, see also methods). To test for a function independent of *GUN5*, *raa20* was crossed with WT Col-0 to obtain homozygous mutants carrying the *ILP1 SNP* (*ilp1^SNP^*) without the *gun5-1* allele (Fig. S1E). We further obtained two allelic knockout (ko) mutants (*ilp1-1* and *ilp1-2*) for *ILP1* in the Col-0 background from publicly available T-DNA insertion collections (Fig. S1F, Yoshizumi et al., 2006). Compared to WT plants grown under standard conditions, the *ilp1* mutants showed a similar rosette size but less pronounced curling of the leaves (Fig. 1A). We also observed the same delayed flowering for *ilp1^SNP^*and the *ilp1* ko plants. For further analysis and to reveal the direct function of ILP1, complementation lines in the *ilp1-1* background were established. To this end, C-terminally HA-tagged ILP1 driven by 35S promoter (*p35S*) or the genomic fragment under the native *ILP1* promoter (*pILP1*) were introduced into *ilp1-1*. While the *p35S::ILP1* lines showed a strong transgene overexpression (∼50-100 fold compared to WT), introducing the *pILP1::ILP1* construct resulted in a moderately elevated *ILP1* mRNA level (Fig. 1C-D). Within the first seven days after germination, *ilp1^SNP^* and *ilp1* ko mutants exhibited a short hypocotyl during de-etiolation and shorter roots when germinated in day-night cycles compared to WT (Fig. 1E-F, Yoshizumi et al., 2006) The perturbed growth of *ILP1*-deficient plants was fully rescued to WT levels in the *p35S::ILP1* and *pILP1::ILP1* complementation lines (Fig. 1E-F). In summary, the selected *ilp1^SNP^* mutant resembled *ilp1* ko mutants in all aspects tested, indicating that this mutant lacks a functional *ILP1*.

**Figure 1:**
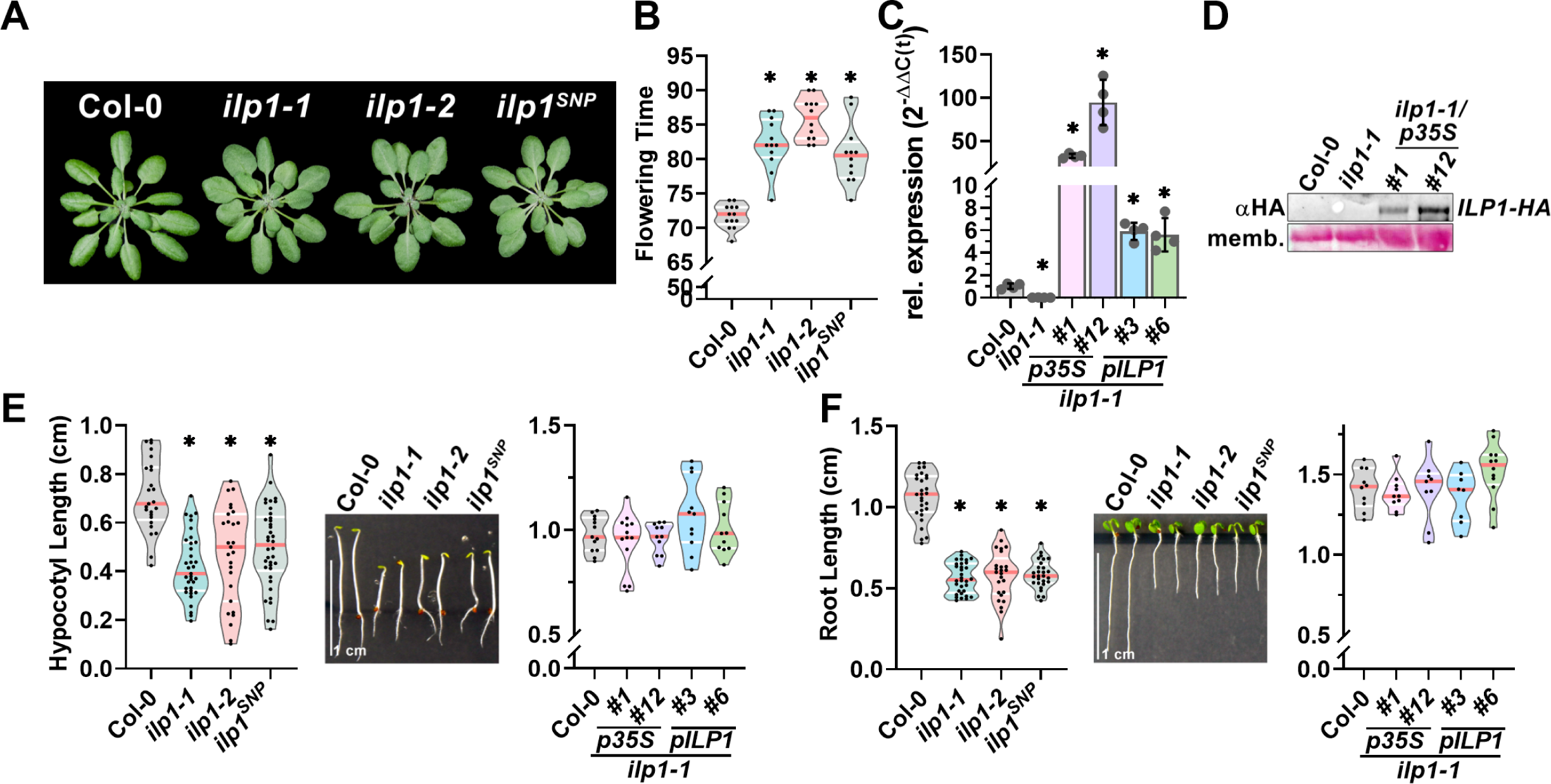
Phenotypes of *ilp1^SNP^*resemble *ilp1* knockout mutants while overexpression (OE) and genetic complementation lines display Col-0-like phenotype. (**A**) Rosette phenotype of 6-week old Col-0, *ilp1-1*, *ilp1-2,* and *ilp1^SNP^* grown under short-day conditions. (**B**) Flowering time (represented as number of days to bolting) of Col-0, *ilp1-1*, *ilp1-2,* and *ilp1^SNP^* (n>10). The red lines inside the violins show the median of values and the white lines the quartiles. (**C**) mRNA expressions of *ILP1* in 4-week-old OE and complementation lines relative to Col-0 at t0 and *SAND* was used as reference gene (n=4). Data are mean ± SD. Asterisks indicate statistical significance compared to Col-0 by Student’s *t*-test (\**p<*0.05). (**D**) Protein levels of ILP1-HA in 4-week-old Col-0, *ilp1-1,* and OE lines. (**E**) Hypocotyl lengths of 4-day-old dark grown Col-0, *ilp1-1*, *ilp1-2, ilp1^SNP^* (n≥25) and OE and complementation lines (n≥11). Asterisks indicate statistical significance compared to Col-0 by Student’s *t*-test (\**p<*0.05). The red lines inside the violins show the median of values and the white lines the quartiles. (**F**) Root lengths of 6-day (n≥25) and 7-day-old (n≥8) Col-0, *ilp1-1*, *ilp1-2, ilp1^SNP^* and *ilp1-1* OE and complementation lines under short-day (SD) conditions, respectively. Asterisks indicate statistical significance compared to Col-0 by Student’s *t*-test (\**p<*0.05). The red lines inside the violins show the median of values and the white lines the quartiles.

### Perturbation of ILP1 and other components of the spliceosome results in a more pronounced activation of anthocyanin biosynthesis in HL

Next, we analysed the HL-mediated activation of anthocyanin biosynthesis in *ILP1*-deficient plants (Fig. 2). After HL exposure, *ilp1* mutants showed higher levels of anthocyanins (Fig. 2A-B), which were reduced to WT-level in *ilp1-1* complementation lines (Fig. 2C). Transcript abundance for the transcription factor *PAP1* and the biosynthesis gene *DFR* rose more quickly and to a higher level in *ilp1-1* after the HL shift. The HL induction of pathway genes resulted in a faster accumulation of anthocyanins which exceeded the level observed for WT plants during a time-resolved kinetic analysis (Fig. 2D-E). Also, after cold treatment (4°C) for seven days in short-day conditions, *ILP1*-deficient plants showed higher anthocyanin accumulation than WT plants (Fig. S2). ILP1 was reported to interact with NTR1, another component of the spliceosomal complex, and *ilp1* and *ntr1* mutants showed similar growth and developmental phenotypes to each other (Yoshizumi et al., 2006; Wang et al., 2019). Here, we confirmed a similarly altered hypocotyl and root growth in both *ilp1-1* and *ntr1-1* (Fig. 2G-H). it is notable that also NTR1 deficiency resulted in elevated anthocyanin levels after HL exposure, and similar contents were detected in *ntr1-1* and *ilp1-1* (Fig. 2I). From the forward genetic screen, two additional allelic mutants, *raa7* and *14,* were isolated (Fig. S1B) and whole genome sequencing revealed that both mutants contain the same SNP in *PLEIOTROPIC REGULATORY LOCUS 1* (*PRL1*, chromosome 4: 9026190, C>T, Supplemental Table S7). PRL1 represents another component of the spliceosomal complex (Koncz et al., 2012), and the *prl1-2* T-DNA insertion mutant resembled *ilp1* and *ntr1* mutants in having a short hypocotyl during etiolation, a short primary root and higher anthocyanin accumulation in HL compared to WT plants (Fig. S3A-F).

**Figure 2:**
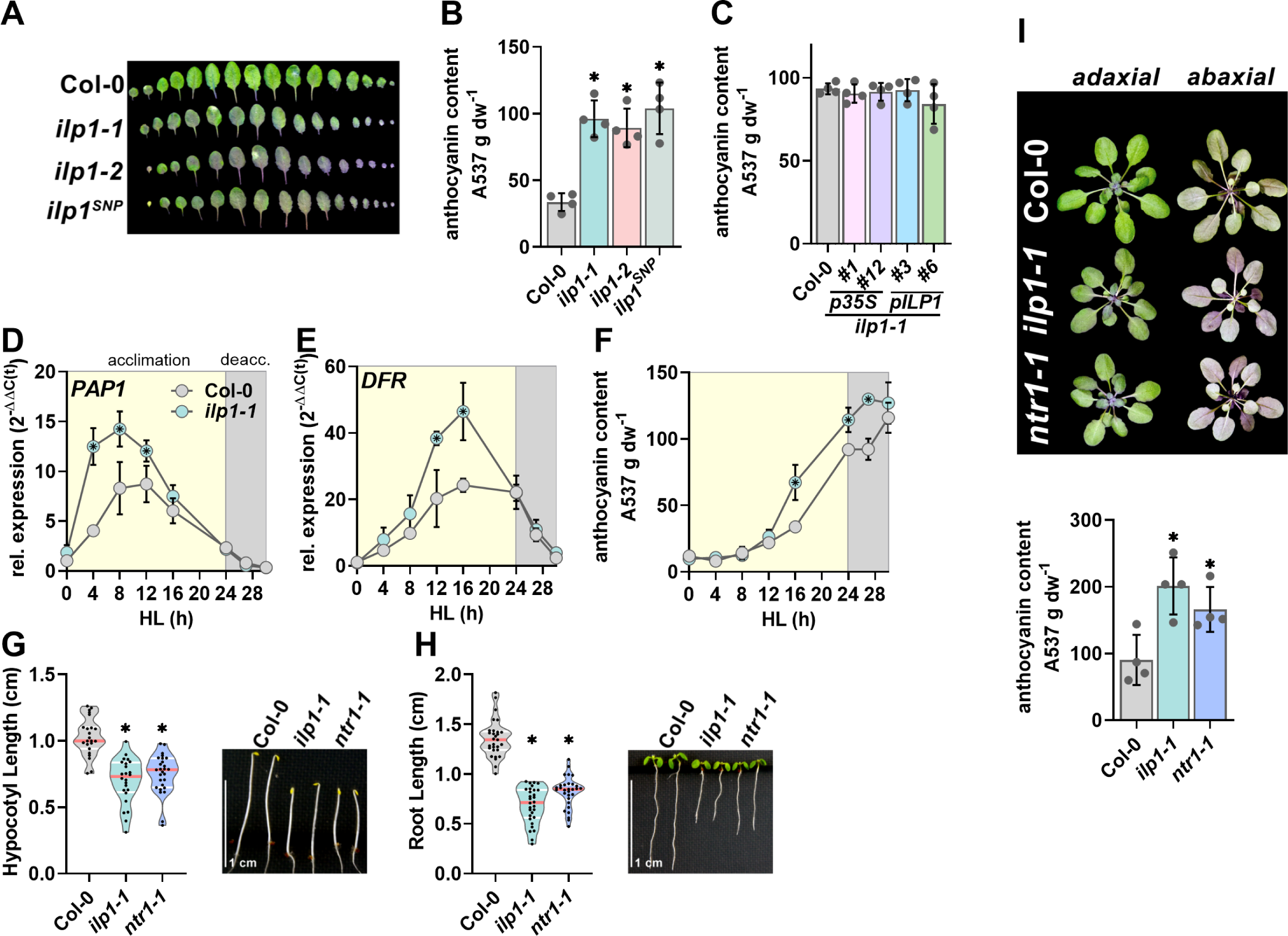
Mutants of *ILP1* and *NTR1* of the ILS have similar phenotypes and response to HL. (**A-C**) Leaf phenotype (**A**) and anthocyanin contents (**B**) of the three mutant alleles of *ILP1* and anthocyanin contents of OE and genetic complementation lines (**C**) after 24 h of HL exposure. (**D-F**) Kinetic analysis of *PAP1* (D) and *DFR* (E) expression, and (F) anthocyanin accumulation during the course of HL acclimation and de-acclimation in 4-week old Col-0 and *ilp1-1* mutants. Transcripts were compared relative to Col-0 at t0 and *SAND* was used as reference gene (n=4). *PAP1, Production of anthocyanin pigment 1; DFR, Dihydroflavonol 4-reductase*. (**G-I**) Hypocotyl length (n≥23, G), root length (n≥29, H), and (I) anthocyanin accumulation phenotype (n=4) of *ntr1-1* and *ilp1-1*. Data are mean ± SD. Asterisks indicate statistical significance compared to Col-0 by Student’s *t*-test (\**p<*0.05). The red lines inside the violins show the median of values and the white lines the quartiles.

Previous analysis revealed a function of ILP1 and NTR1 for transcriptional elongation of *MIR* genes encoding miRNAs (Wang et al., 2019), and miRNA-dependent regulation of anthocyanin biosynthesis has been reported (e.g., Gou et al., 2011; Wang et al., 2016; He et al., 2019; Wang et al., 2020). Therefore, we tested the *serrate* (*se1*) mutant (Lobbes et al., 2006; Laubinger et al., 2008) severely affected in miRNA biogenesis for the activation of anthocyanin biosynthesis during HL acclimation (Fig. S3). Compared to WT, leaves of the *se1* mutant showed a serrated phenotype, confirming previous results. However, we detected WT-like contents of anthocyanins (Fig. S3G) after 24 h and stimulation of flavonoid biosynthesis transcription factors and pathway genes after short-term HL exposure (Fig. S3H) in the *se1* mutant. These results indicate that SE1-dependent miRNA biogenesis plays a minor role in anthocyanin biosynthesis activation in the conditions tested here. However, we cannot exclude a role for miRNAs that are generated via other routes independent of SE1.

We concluded that ILP1, NTR1 and PRL1 exhibit a repressive function on anthocyanin accumulation, and knockout of these factors leads to a pronounced activation of the pathway under acclimation-relevant conditions.

### Conditional and constitutive perturbation of gene expression in *ilp1-1*

The more pronounced accumulation of anthocyanins and deregulation of anthocyanin biosynthesis in HL prompted us to investigate the global transcriptional response of Col-0 and *ilp1-1* to HL (Fig. 3). We performed RNA sequencing analysis (RNAseq) with samples harvested at different time points of HL exposure. Samples were harvested at t0 (2 h after the onset of light), 4, 8, and 16 h of HL treatment, as well as after 24 h of HL exposure followed by six hours of de-acclimation in normal light conditions (30 hours in total). Principal component analysis (PCA) of the biological replicates of each time point indicated high consistency across the replicates (Fig. 3A). We observed distinct trajectories of changes in the transcriptomes of both Col-0 and *ilp1-1*, which were influenced by the duration of light exposure (PC1) and genotype differences (PC2). Overall, both genotypes showed an acclimation/de-acclimation response with pronounced gene expression changes after 4-16 h and a (partial) reversal of the changes during de-acclimation (30 h), indicating that HL acclimation was still functional to some extent in *ilp1-1* (Fig. 3A). The transcript changes were also influenced by the circadian clock which was shown to be influenced by ILP1 function. However, z-transformed FPKM (fragments per kilobase of transcript per million mapped reads) values plotted as a heatmap revealed pronounced differences in *ilp1-1* compared to the WT samples, particularly in the first hours of the HL treatment and of the recovery phase (Fig. 3B). Transcripts of 20,545 genes in common were identified across all samples, and were grouped into four major clusters (I-IV) based on their expression profiles. These clusters represented transcripts induced or repressed during acclimation and showed a reversal of the response during de-acclimation (clusters I+II+IV) (Fig. 3B). Additionally, transcripts showing continuing repression (cluster II) and induction during HL exposure (part of cluster II+IV) were identified. Comparison of *ilp1-1* and WT Col-0 revealed significantly differentially expressed genes (DEGs) between the genotypes in all clusters. For instance, transcripts for the regulation and biosynthesis of flavonoids showed an induction in Col-0 but a more pronounced induction in *ilp1-1* in the first hours of HL exposure (Fig. 3C). For instance, transcripts for the regulation and biosynthesis of flavonoids showed an induction in Col-0 but a more pronounced induction in *ilp1-1* in the first hours of HL exposure (Fig. 3C). Likewise, while repressed in WT after the shift to HL, some transcripts involved in auxin biosynthesis and signalling were expressed at a higher level in *ilp1-1* (Fig. 3C, Supplemental Tables S1 and S2).

**Figure 3:**
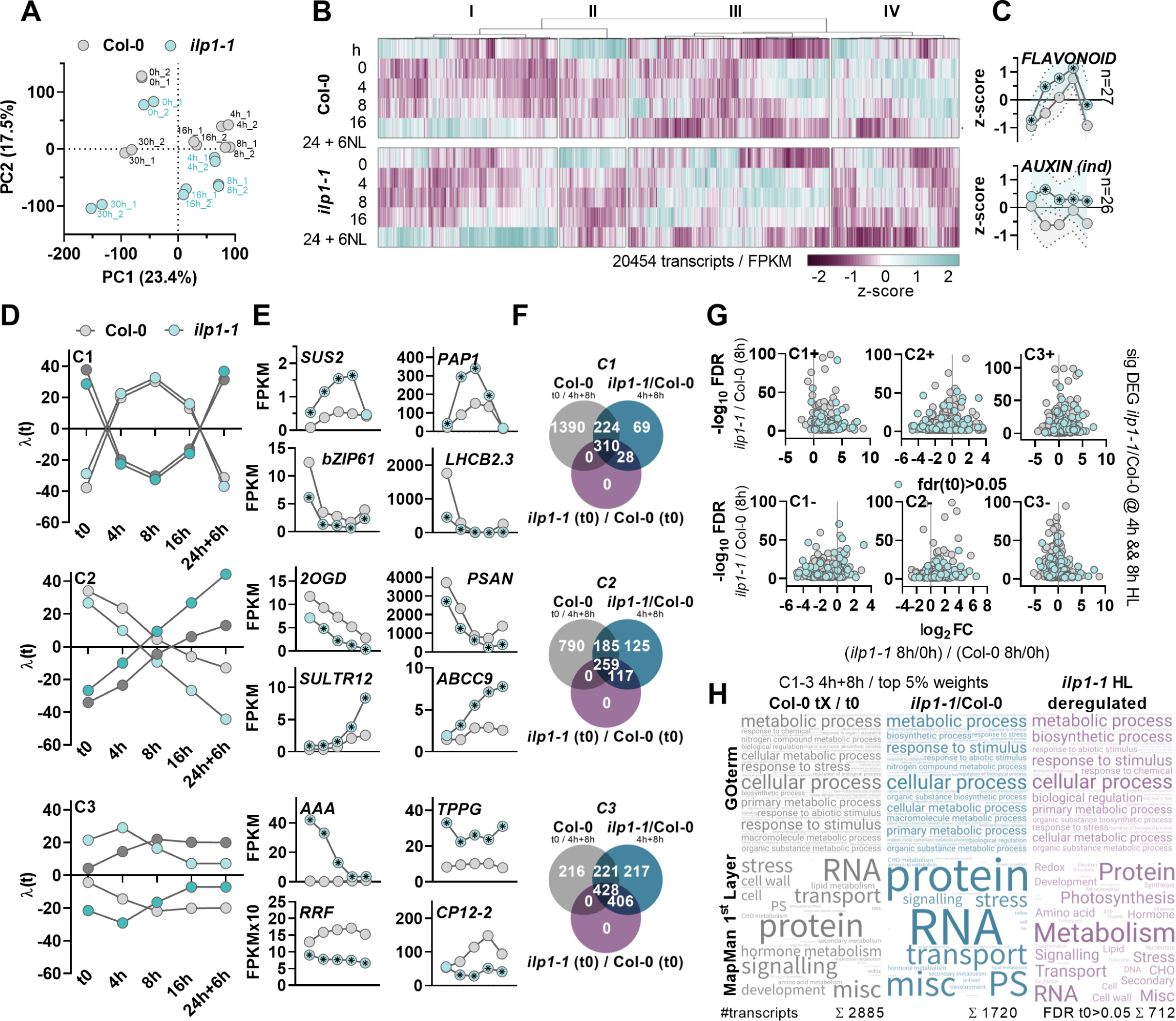
Global transcriptome and TMEA analysis revealed constitutive and conditional perturbation of gene expression in *ILP1*-deficient plants. (**A**) PCA plot showing the scattering of replicate samples (1 and 2) of Col-0 and *ilp1-1* treated with HL for the indicated time points. Plants analysed after 30 h were harvested after 6 h recovery in NL following 24 h HL treatment. (**B**) Heatmap summarising expression changes of 20. 454 transcripts identified at all time points throughout the HL kinetic (FPKM>0). FPKM values were z-transformed, and colours indicate changes in the z-score. (**C**) Mean z-transformed FPKM values for transcripts encoding proteins involved in the regulation and biosynthesis of flavonoids (top), and biosynthesis of and response to auxin (bottom). The transcripts were selected based on MapMan bins and fdr<0.05 at 4 h and 8 h HL. Only transcripts induced (ind) in *ilp1-1* are depicted for the auxin bin. Asterisk inside symbols indicates significant differences (p<0.05) from a pairwise *t-*test between Col-0 and *ilp1-1* samples at each time point. Shades indicate standard deviation. *FLAVONOID* (n=27) and *AUXIN(ind)* (n=26) transcripts. (**D**) Time course of the constraint (C) potentials (λ(t)) for C1, C2, and C3 obtained from TMEA analysis. Bright colours indicate the negative (-), and dark shades the constraints’ positive (+) trajectory. (**E**) Exemplary transcripts representing C1-C3. Transcript amounts are given as log(fpkm+1). Asterisks inside the circles indicate significant differences determined DeSeq2 using Col-0 and *ilp1-1* samples of the HL kinetic (fdr<0.05). *FLAVONOID* (n=27) and *AUXIN(ind)*, n=26 transcripts. *SUS2*, *Sucrose synthase 2. PAP1, Production of Anthocyanin Pigment 1. bZIP61, Basic-leucine zipper (BZIP) transcription factor 61. LHCB2.3, Light-harvesting complex protein B2.3. PSAN, Photosystem I reaction centre subunit PSI-N. 2OGD, 2-oxoglutarate (2OG) and Fe(II)-dependent oxygenase superfamily protein (AT3G46490). SULTR12, Sulfate transporter 12. ABCC9, ABC-type xenobiotic transporter. AAA, AAA-ATPase AT2G18193. TPPG, Probable trehalose-phosphate phosphatase G (TPP6). CP12-2, Calvin cycle protein CP12-2. RRF, Ribosome-releasing factor*. (**F**) Venn diagrams depicting common and differentially expressed genes (DEG, DeSeq2, fdr<0.05) of the following comparisons: 1. Col-0 DEG 4h and 8h relative to t0 (grey). 2. DEGs in *ilp1-1* relative to Col-0 after 4 h and 8 h HL treatment (blue). 3. DEG in *ilp 1-1*relative to Col-0 at t0 (purple). The analysis was limited to the top and lowest 5 % of transcripts (weights) of C1-3. (**G**) Volcano plots depicting DEG of C1-3 (top 5% weights for the positive and negative C) showing an induction or repression upon HL exposure of Col-0 and *ilp1-1*. Firstly, relative expression change between t0 and 8 h HL was calculated for each genotype individually (based on log(fpkm+1), genotype 8h/t0)). The ratio between *ilp1-1* (8h/0h) / Col-0 (8h/0h) revealed relative differences in the induction and repression of transcripts abundance in HL. Blue symbols indicate transcripts exclusively deregulated in *ilp1-1* in HL (ilp1-1 (t0) / Col-0 (t0), fdr>0.05). FC, fold change. (**H**) Word clouds showing the Gene ontology (GO) term (top row) and MapMan bin enrichment among DEG of (1) Col-0 4 h and 8 h / t0 (grey), (2) *ilp1-1* / Col-0 at 4 h and 8 h(blue), (3) exclusively HL deregulated transcripts in *ilp1-1*. For (1) and (2), transcripts represented by C1-C3 of TMEA analysis (top 5% weights for the positive and negative C) and DEG at 4 h and 8 h (fdr<0.05) of the HL kinetics were used. For (3) transcripts differentially expressed in Col-0 at 4 h and 8 h of HL relative to t0, in *ilp1-1* / Col-0 (4 h and 8 h HL) but not in *ilp1-1*/Col-0 at t0 (fdr>0.05) were analysed (overlap of grey and blue and exclusively blue in (F)). (**G**) Anthocyanin contents of 4-week old Col-0, *ilp1-*1, and *se1* after 24 h of HL exposure. Data are mean ± SD. Asterisks indicate statistical significance compared to Col-0 by Student’s *t*-test (\**p<*0.05). (**H**) Relative expressions of *CHS*, *FLS1*, *PAP1*, and *DFR* in 4-week old Col-0, *ilp1-1*, and *se1* before and 8 h after HL exposure. Transcripts were compared relative to Col-0 at t0 and *SAND* was used as reference gene (n=4). Data are mean ± SD. Asterisks indicate statistical significance compared to Col-0 by Student’s *t*-test (\**p<*0.05). CHS, Chalcone synthase; FLS1, Flavonol synthase 1.

To allow for a comprehensive description of the HL acclimation response and to reveal commonalities and differences between Col-0 and *ilp1-1,* a thermodynamically <u>m</u>otivated <u>e</u>nrichment <u>a</u>nalysis (TMEA, Schneider et al., 2020) was conducted (Fig. 3D and Supplemental Table S1,). TMEA is an energy-based method employing a surprisal analysis to allow the mathematical description of the contribution of individual transcripts (the weights) to changes of a biological steady state (here, the transcriptome). The changes imposed on a system are described by ’constraints’ (C) which represent different responses of transcripts and are tested for functional enrichment by TMEA. Constraints describe the behaviours of the transcriptome response over time. Each transcript has weights associated with each constraint describing its relevance. Consequently, transcripts possessing high weights to a constraint share common expression states during the experiment. For example, transcripts exhibit a typical pattern of acclimation/de-acclimation, with some being induced and others repressed during exposure to HL and the opposite occurring during the recovery phase (C1 positive or negative, Figs. 3D and S4A-B). Others are increasingly induced or repressed during HL, without returning to their original expression level during de-acclimation (mainly represented by C2, Fig. S4C-D). The C3 of the TMEA analysis represents transcripts with distinct and constitutive expression changes in *ilp1-1* compared to Col-0 during the HL kinetic (Fig. S4E-F). For further analysis, transcripts with the highest contribution to the positive and negative C1-3 were considered (top 5% of weights), and exemplary transcripts for each constraint are provided in Fig. 3E (Supplemental Table S1). For example, *SUCROSE SYNTHASE 2 (SUS2)* and *PAP1 (MYB75)* showed the same trajectory in both genotypes. However, they were massively overexpressed in *ilp1-1* upon HL shift corroborating the changed regulation of anthocyanin biosynthesis (see Figs. 2 and 3C) and revealing altered response of transcripts for carbohydrate metabolism (see C2). In contrast, *PSAN* (*PHOTOSYSTEM I REACTION CENTER SUBUNIT N*) and *SULTR1;2* (*SULFATE TRANSPORTER 1;2*) showed stronger repression during HL and induction during recovery, respectively, in *ilp1-1*. For others, such as a *AAA-ATPase* (*AT2G18193*), the expression did not change in Col-0 during HL but was strongly repressed in *ilp1-1* starting from a high expression level at t0 or was induced in Col-0 but not HL responsive in *ilp1-1* (e.g., *CALVIN CYCLE PROTEIN 12-2*, *CP12-2*) (see C3). C1 is enriched with transcripts grouped into the GO-terms “anthocyanin/flavonoid biosynthesis”, “rRNA biogenesis”, “ribosomal maturation”, “translation”, “carbohydrate metabolism”, “response to biotic stimuli” (Fig. S4A-B), all of which were previously identified as major groups regulated during plant acclimation to various abiotic stresses (Garcia-Molina et al., 2020). We recognised that some mRNAs were differentially expressed in *ilp1-1* relative to Col-0 before the HL exposure (at t0) (Figure S4G). Hence, the observed (absolute) differences in FPKM for some transcripts between the mutant and WT in HL were partially influenced by the existing changes at t0. In other words, although exhibiting changes in (absolute) transcript abundance during HL exposure, these transcripts were similarly induced or repressed as in the WT Col-0, and hence, HL regulation was not influenced by *ILP1*-deficiency. To identify the truly HL de-regulated genes in *ilp1-1*, we compiled a list of transcripts represented by C1-3 and applied the following filter strategy to reveal: (1) DEGs upon HL shift in the WT (DEGs for Col-0 t0/ 4h+8h fdr<0.05, Fig. 3F grey circles), (2) DEGs in *ilp1-1* (DEGs *ilp1-1*/Col-0 at 4h+8h, Fig. 3F blue circles), and (3) DEGs in *ilp1-1* before the HL shift (*ilp1-1* (t0) / Col-0 (t0), Fig. 3F purple circle). The overlaps showed that approximately 50% of DEGs were truly HL de-regulated in *ilp1-1* (Fig. 3F, overlap grey and blue). Of these, approximately 50% were exclusively de-regulated in *ilp1-1* (Fig. 3F, exclusively blue, Table S3). We next plotted the quotient (as log_2_ fold change) of the induction and repression, respectively, of transcripts in Col-0 and *ilp1-1* after 8 h HL relative to t0 against the fdr for the expression change in *ilp1-1*/Col-0 (8h) (Fig. 3G, Supplemental Table S4). In agreement with the trajectory of the identified constraints and the observed relative changes for individual transcripts (Fig 3D-E), DEGs in *ilp1-1* showed stronger induction (positive log_2_ FC) and repression (negative log_2_ FC), respectively, in each constraint. Across the DEGs in *ilp1-1*, those exclusively de-regulated in HL followed the overall trend of expression changes observed in the C (Fig. 3G, blue symbols, *ilp1-1/*Col-0 t(0), fdr > 0.05). GO term and MapMan bin analysis revealed transcripts from the following categories: “cellular response”, “metabolic process”, and “response to stimulus” involved in “RNA”, “protein”, and “hormone metabolism, transport and signalling” as highly enriched in C1-3 in HL treated WT (Fig. 3H, grey and Supplemental Table S5). It is noteworthy that the same GO terms were enriched across the DEGs in *ilp1-1* compared to WT with a pronounced enrichment of transcripts for “RNA”, “protein “, and “photosynthesis” at 4 and 8 h of HL exposure (Fig. 3H, blue, Fig. S4G, and Supplemental Table S5). Considering the DEGs de-regulated in *ilp1-1* upon HL shift without being DEG at t0 (Fig 3F, overlap grey/blue and exclusively blue), GO terms also enriched in the WT were found (Fig. 3H, purple and Supplemental Table S5). However, based on MapMan bins, gene products involved in “protein”, “primary and secondary”, “amino acid”, “carbohydrate”, “lipid”, “hormone”, “cell wall”, “metabolism”, “transport” and “signalling” were enriched among the HL-deregulated DEGs in *ilp1-1*. In summary, *ILP1* is a crucial factor for the proper HL acclimation and the de-acclimation response of plants, and deficiency provokes a pronounced alteration of gene expression, particularly affecting genes essential for plant metabolism.

### Analysis of splicing events in WT and *ilp1-1*

Splicing of introns is one crucial part of the maturation of transcripts. Among others, altered or regulated splicing results in retention of introns (RI), skipping of exons (SE), or alternative 5’ (A5) and 3’ (A3) sites, thereby allowing accumulation of functional mRNAs or alternative transcripts resulting in the expression of protein isoforms. The spliceosomal component ILP1 affects these splicing events, and *ilp1-1* mutants showed an apparent splicing defect for RI, SE, A5 and A3 events not only under normal but also under HL conditions (Fig. 4A-C, Table S6). In WT plants, >90% of the detected transcripts showed a low level of RI, a high inclusion level of exons and a varying degree for the presence of A5+A3 (Fig. 4A-C). Also, the global analysis of splicing events in WT plants revealed constant splicing of primary transcripts across the different time points from the HL shift experiment (Fig. 4A-C and S5A), suggesting that dynamic splicing did not underpin the HL-acclimation response of WT plants. In contrast, we detected significant alteration of RI and SE events in *ILP1*-deficient plants (Fig. 4A-B and S5A). The *ilp1-1* mutant showed a significantly increased inclusion level for RI (approximately +20% compared to WT) and a lower inclusion level for SE (Fig. 4A-C and S5A). is It is noteworthy that, compared to the constant inclusion level in WT, a dynamic adjustment of splicing was observed in *ilp1-1* during HL acclimation, and after 6 h of recovery from 24 h HL exposure inclusion level in *ilp1-1* was more similar to WT than before the treatment (Fig. 4A-C and S5A). These results suggest that ILP1 is vital for the proper splicing of transcripts, but splicing can still occur, and ILP1-independent splicing is induced during HL acclimation in the absence of ILP1. GO-term and MapMan bin analysis of differentially spliced transcripts across the HL time course in *ilp1-1* revealed a pronounced enrichment of mRNAs involved in “Metabolic Processes” such as nucleotide, amino acid, carbohydrate, lipid, hormone and secondary metabolism (Fig 4D-E). Although the inclusion level did not significantly change globally, we observed individual changes in splicing events in WT plants and selected transcripts which showed alternative splicing during the HL time course using ANOVA analysis (adjusted p-value<0.05). Next, we asked whether the adjustment of splicing/ inclusion level depends on ILP1. To this end, differentially spliced transcripts identified in WT were tested for alternative splicing in *ilp1-1.* For this, the adjusted p-values of the ANOVA for WT and *ilp1-1* were plotted (Fig. 4F). Remarkably, more than 80% of the splicing events were ILP1-dependent suggesting that indeed ILP1 mediated (alternative) splicing contributes to the HL acclimation response of plants (Fig. 4F). GO term analysis showed that transcripts encoding enzymes and proteins for various branches of plant metabolism were substantially affected in *ilp1-1* (Fig. 4G).

**Figure 4:**
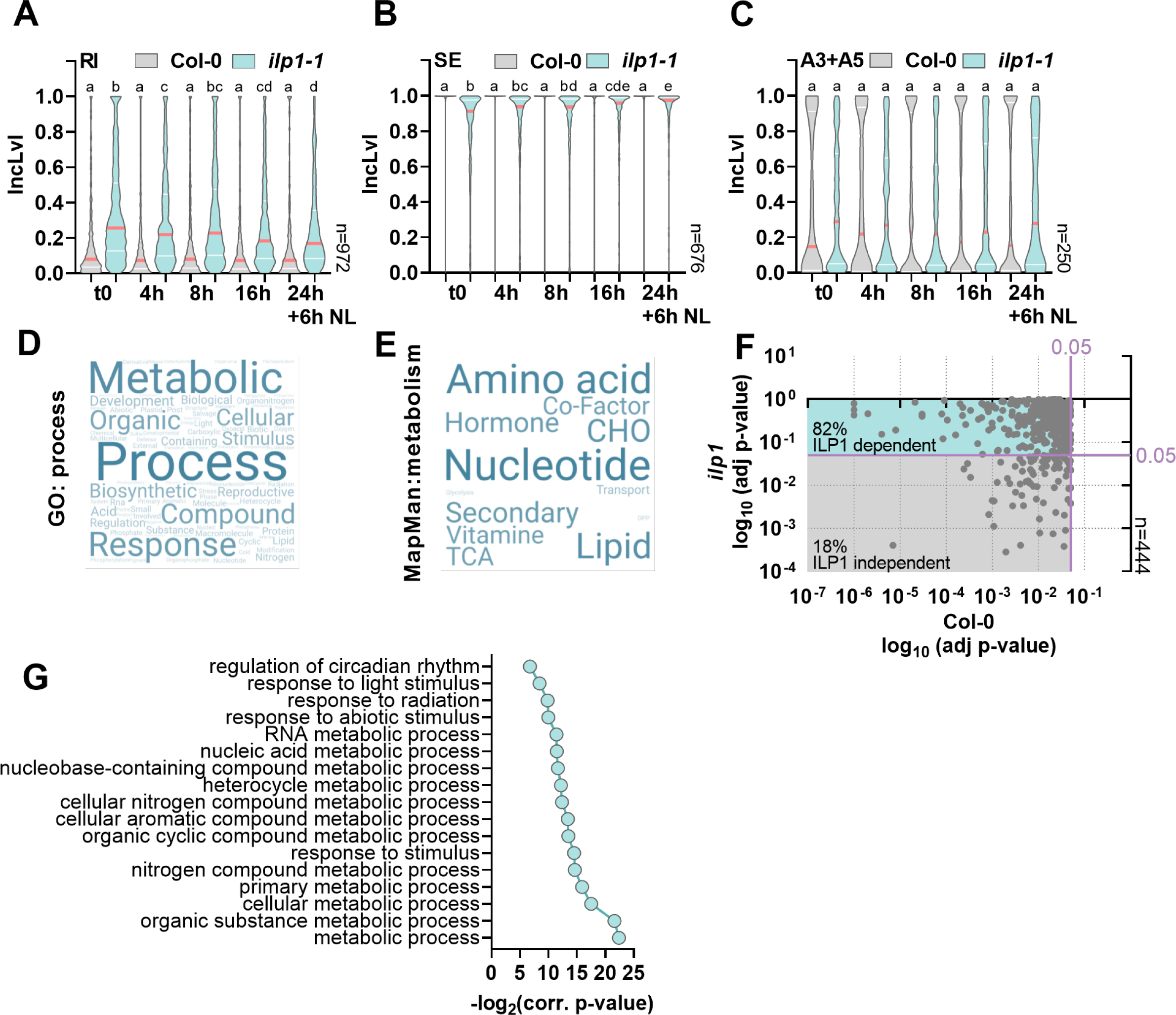
Global analysis of splicing events in Col-0 and *ilp1-1* during HL treatment. (**A-C**) Violin plots depicting changes in inclusion levels (IncLvl) for splicing events in Col-0 and *ilp1-1*: (A) Retained Intron (RI), (B) Skipped Exon (SE) and Alternative 3’ and 5’ ends (A3+A5). The red lines inside the violins show the median IncLvl and the white lines the quartiles. Transcripts showing a global change in IncLvl between Col-0 and *ilp1-1* across all time points were analysed (two-way ANOVA, fdr<0.05). Letters above the violins indicate significance groups identified with one-way ANOVA analysis (Tukey’s multiple comparison tests, adj p<0.05). (**D**) Word cloud showing GO-term enrichment of transcripts identified in (A-C). (**E**) Word cloud showing MapMan enrichment of differentially spliced transcripts in *ilp1-1* encoding gene products involved in metabolic pathways (from the analysis shown in A-C). The list of MapMan bins enriched in the data set was filtered for the keyword ’metabolism’. (**F**) Identification of transcripts with ILP1-dependent differential splicing during HL (RI+SE+A3+A5). The list of differentially spliced transcripts (DST) in Col-0 and *ilp1-1* samples were filtered for global changes of IncLvl in Col-0 (one-way ANOVA, adj p-value <0.05). The adj p-values for Col-0 and *ilp1-1* of the transcripts were plotted against each other. The blue area highlights transcripts not differentially spliced in the mutant but in the COL-0 (*ilp1-1* adj P>0.05 and Col-0 adj p<0.05). The grey area indicates DST in Col-0 and *ilp1-1* (adj p<0.05) and are thus ILP-independent. The purple lines depict the significance threshold (adj p<0.05). (**G**) GO-term enrichment of DST for the ILP1-dependent splicing events identified in (F). GO term enrichment analysis was performed using https://go.princeton.edu/cgi-bin/GOTermFinder.

Splicing may affect the stability of mRNAs and, eventually, the expression of regulatory factors for HL acclimation. We sought to understand the connection between splicing and mRNA contents during HL, mainly focussing on ILP1-dependent effects. Surprisingly, no global correlation between inclusion and mRNA levels was observed for *ilp1-1* or WT plants (Fig. S5B), suggesting that differential splicing overall did not affect transcript stability but may contribute to the accumulation of alternative protein species during HL acclimation.

Taken together, ILP1-deficiency resulted in constitutive and conditional changes of splicing affecting transcripts for plant primary and secondary metabolism. Of note, although the known genes for enzymes and transcriptional regulators of anthocyanin biosynthesis were de-regulated, no apparent candidate gene functioning upstream of anthocyanin biosynthesis regulators could be identified from the transcriptome and splicing analysis in *ilp1-1* that would explain the changed anthocyanin accumulation during HL acclimation.

### Metabolic analysis revealed conditional alteration of carbon and nitrogen metabolism in *ilp1-1*

The pronounced perturbation of gene expression and altered splicing in *ilp1-1,* particularly of mRNAs encoding gene products crucial for various branches of plant primary and specialised metabolism during the HL acclimation, prompted a detailed analysis of different metabolites. First, we focused on carbohydrate and carbon metabolism. We found that *ILP1*-deficiency resulted in a significantly elevated level of starch at the end of the night (EoN), 2 h after the onset of light (t0) and long-term (24 h) HL exposure (Fig. 5A-B). Also, short-term HL treatment caused an almost 2-fold increase in sucrose (Fig. 5C), an intermediate of sucrose biosynthesis (sucrose 6^F^-phosphate), and the regulatory sugar-phosphate trehalose 6-phosphate in *ilp1-1* relative to Col-0 (Fig. 5D). Importantly, changes in the sugar contents relative to WT were only observed upon HL treatment of the mutant. Significantly higher starch accumulation was also observed during a normal day-night cycle in short-day conditions in tissues of *ilp1-1* compared to WT plants (Fig. S6A). While the starch degradation at night was similar to WT, net accumulation of starch showed some sign of being elevated during the day in the absence of ILP1, but the differences between mutant and WT were not statistically different (Fig. S6B). Starch and carbohydrate accumulation strictly depend on photosynthesis and carbon fixation. At t0 or under normal light conditions, photosynthetic parameters, such as Fv/Fm, qP, NPQ, and carbon (C) fixation, were WT-like in *ilp1-1* (Fig. S6C-D). However, after 8 h and 24 h HL exposure or with increasing light intensities even exceeding 500 µmol photons m^-2^ s^-1^, a striking difference between *ilp1-1* and Col-0 plants was observed. While the photosynthetic performance of WT plants first increased (8 h) and then declined until 24 h HL exposure, *ilp1-1* mutants showed higher values for Fv/Fm, qP, NPQ, Y(II) and Y(NPQ) and lower values for non-regulated energy dissipation (Y(NO)) after 24 h of HL treatment (Fig. S6C). Also, *ilp1-1* was characterised by a higher CO_2_ fixation rate than WT in a light-response curve with plants from t0 and after 24 h HL (Fig. S6 D). Chlorophyll contents and steady-state level of photosynthetic proteins were WT-like in *ilp1-1* during HL treatment (Fig. S6E-F). Likewise, marker genes for reactive oxygen species (ROS) stress were overall expressed at WT-like or even lower levels in *ilp1-1* (Fig. S7A), and ROS staining did not indicate altered accumulation of ROS upon HL treatment in the allelic *ILP1* mutants (Fig. S7B-C). Also, a targeted RNAseq data analysis revealed WT-like expression of gene products involved in phytohormone biosynthesis and signalling in *ilp1-1* throughout the HL kinetic (Fig. S7D).

**Figure 5:**
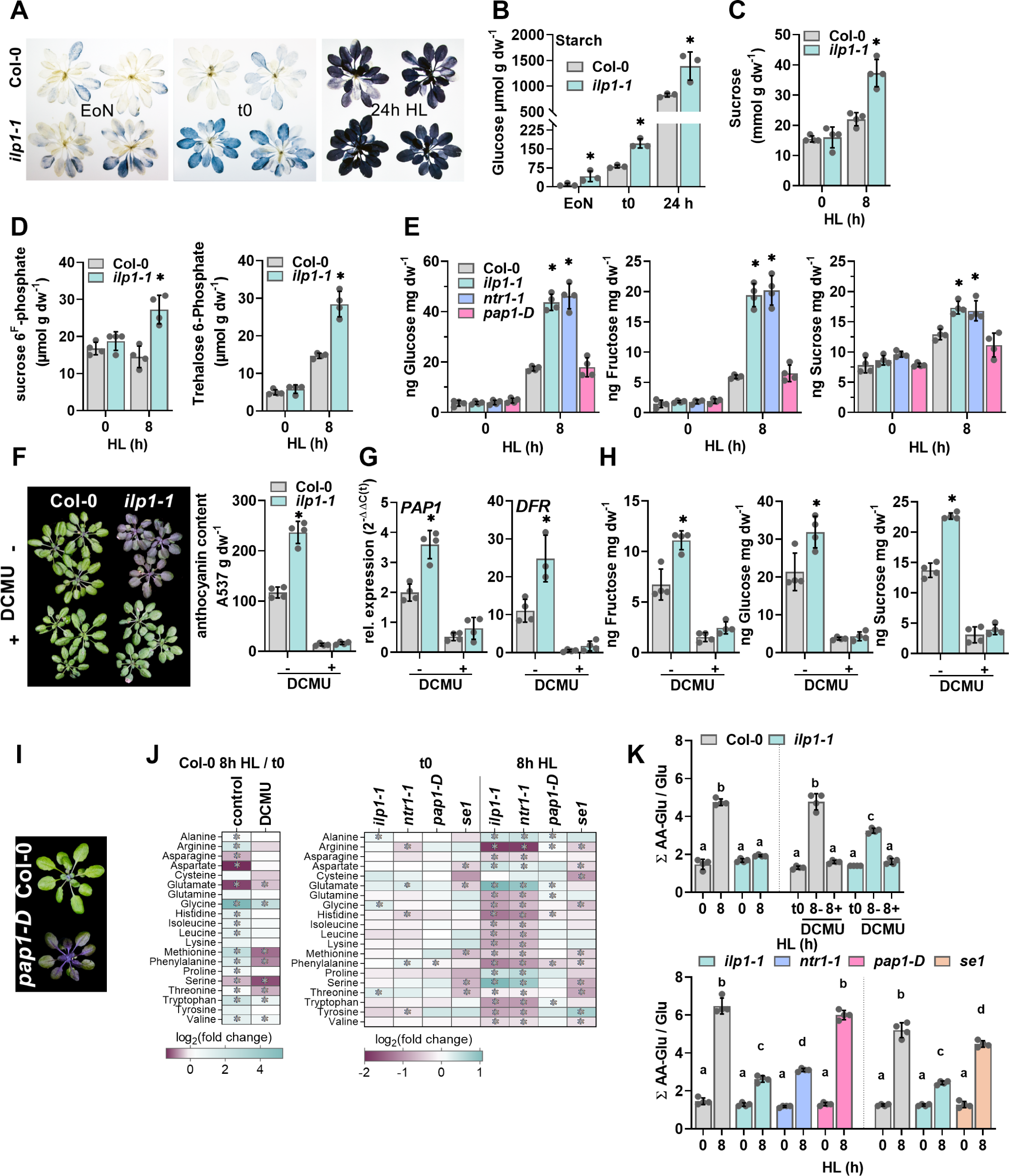
Mutants of the spliceosomal complex, *ilp1-1* and *ntr1-1*, show conditional imbalance of the C:N ratio during HL acclimation. (**A** and **B**) Lugol’s staining (A) and starch contents (B, expressed as glucose equivalents) of 4-week old Col-0 and *ilp1-1* at EoN, before (t0) HL, and 24 h after HL. (**C**-**E**) Content of sucrose (C) and phosphorylated sugars (D) in 4-week old Col-0 and *ilp1-1* before and 8 h after HL. (E) Glucose, Fructose and Sucrose contents in *ilp1-1* and *ntr1-1* display similar levels during HL. (**F**-**H**) Phenotype and anthocyanin contents (F), relative expressions of *PAP1* and *DFR* after 8 h HL (G), and (H) neutral sugar contents in DCMU-treated and untreated plants after 24 h HL. For B-H, Data are mean ± SD (n≥3). Asterisks indicate statistical significance compared to Col-0 by Student’s *t*-test (\**p*<0.05). (**I**) Phenotype of *pap1-D* (dominant *PAP1* overexpression allele) compared to Col-0 after 24 h HL exposure. (**J**) Log_2_ fold changes of the different amino acids compared to the control without DCMU (left panel) or Col-0 (right panel) at the indicated time points. Data are mean ± SD (n=4). Asterisks indicate statistical significance compared to Col-0 by Student’s *t*-test (\**p<*0.05). (**K**) (∑Amino Acids – Glu)/Glu in DCMU-treated plants, spliceosomal mutants, *pap1-D,* and *se1* before and 8 h after HL. Different letters indicate significance groups at p<0.05 by Šidák’s and Tukey’s multiple comparison tests, respectively, as determined by two-way ANOVA.

Besides the ILP1-deficient plants, *ntr1-1* and *prl1-2* mutants also exhibited HL-induced accumulation of glucose, fructose and sucrose beyond WT levels while overaccumulating anthocyanins (Fig. 5E and S3D). To exclude the effects on sugar contents arising from induced anthocyanin biosynthesis, we analysed a dominant *PAP1* overexpression line (*pap1-D*) with marked anthocyanin accumulation (Fig. 5I). Overexpression of *PAP1* and substantial anthocyanin accumulation in *pap1-D* resulted in WT-like levels of soluble sugars before and after short-term HL exposure (Fig. 5E), indicating that the increased carbohydrate contents in *ilp1-1*, *ntr1-1,* and *prl1-2* were not caused by anthocyanin overaccumulation. Instead, DCMU (3-(3,4-dichlorophenyl)-1,1-dimethylurea) application, diminishing photosynthetic electron transfer and carbon fixation, suppressed anthocyanin accumulation in WT and *ilp1-1* in HL (Fig. 5F) and prevented HL-mediated induction of anthocyanin biosynthesis genes (Fig. 5G). After DCMU application, sugar contents were markedly reduced in both genotypes (Fig. 5H), and we found a strong correlation with the (in)activation of anthocyanin biosynthesis in WT and *ilp1-1,* corroborating a direct impact of photosynthetic activity and cellular sugar contents on HL activation of anthocyanin biosynthesis (Zirngibl et al., 2023). The transcriptome analysis indicated a perturbation of metabolism of N-containing metabolites, such as nucleotides and amino acids (see above) and, therefore, amino acid (AA) contents as major sinks for N were analysed utilising LC/MS. Overall, WT-like AA contents in *ilp1-1*, *ntr1-1*, *pap1-D* and *se1* were detected in standard growth conditions (Fig. 5J). In contrast, after HL exposure, a conditional alteration of AA contents became visible in *ilp1-1* and *ntr1-1*. In the WT, Asp, Glu, and Ser contents were reduced while other AAs (e.g., Arg, Gly, Met, Phe, Trp) showed relative enrichment after 8 h HL compared to t0, indicating general stimulation of AA biosynthesis in HL. Remarkably, the contents of most AA were diminished in *ilp1-1*, *ntr1-1* and *prl1-2* after 8 h HL exposure, but Glu levels were markedly increased compared to WT in all three mutants (Fig. 5J and Supplemental Fig. S3E). Although *SE1*-deficiency resulted in changes of some AA after 8 HL, the relative changes compared to WT were overall already present before the treatment (Fig. 5J). The higher anthocyanin accumulation in the *pap1-D* mutant (Fig. 5I) was accompanied by differences in the levels of some AA in the conditions tested compared to WT plants (Fig. 5J). However, in *se1* and *pap1-D,* the alterations of AA pools upon HL acclimation were generally distinct from those observed in *ilp1-1*, *ntr1-1,* and *prl1-2*. Notably, all three spliceosomal complex mutants showed the same alteration of AA metabolism compared to the WT, revealing a role of the spliceosomal complex components for conditional adjustment of N metabolism, particularly AA biosynthesis, during HL acclimation. Glu is one of the most abundant AA in plants and serves as the major amino group donor for the biosynthesis of all AA. Hence, calculating the (∑AA-Glu)/ Glu ratio can indicate changes in AA biosynthesis under HL. In WT plants, an increase of the (∑AA-Glu)/ Glu ratio during short-term HL is indicative of active AA biosynthesis and a decrease in the presence of DCMU, shows a strict dependency on photosynthetic activity (DCMU application, Fig. 5J-K). By contrast, across five independent experiments, *ilp1-1* showed significantly higher Glu levels and a lower (∑AA-Glu)/ Glu ratio compared to the WT Col-0, while this alteration was not observed in standard growth conditions (t0) (Fig. 5K and Supplemental Fig. S3F). The same was observed for *NTR1*- and *PRL1*-deficient plants (Fig. 5K and Supplemental Fig. S3F). Glu also serves as NH_4_^+^ acceptor in the course of N-assimilation by GLUTAMINE (Gln) SYNTHETASE (GS) in the cytosol and plastids. In HL-treated WT plants, the Gln/Glu ratio increased by 4.3-fold (± 0.68 SEM, n= 19, 5 experiments, Supplemental Table S8) relative to the normal light condition. On the contrary, the Gln/Glu ratio of *ilp1-1* only increased by 1.4-fold (± 0.12 SEM, n= 20, 5 experiments, Supplemental Table S8), which was interpreted as a potential limitation in the conditional stimulation of Gln biosynthesis and N-assimilation in HL. Pronounced anthocyanin accumulation in *pap1-D* and perturbation of miRNA biogenesis in *se1* did not affect the (∑AA-Glu)/ Glu ratio (Fig. 5K). In summary, the lack of ILP1, NTR1, and PRL1 resulted in a conditional perturbation of carbon-and N-metabolism, indicating that the spliceosomal components are crucial factors for the adjustment of the C:N balance during HL acclimation.

### *ilp1-1* and *ntr1-1* show symptoms of nitrogen deficiency but unaltered N content

The obtained results pointed to a disturbance of N-metabolism when the ILP1 function is perturbed. Indeed, a targeted analysis revealed that more than 95% of transcripts reported to be regulated under N-limitation (Scheible et al., 2004; Krapp et al., 2011) were also differentially expressed in *ilp1-1* relative to Col-0 (Fig. 6A). During normal light conditions (t0, prior to HL), *ilp1-1* exhibited a similar pattern of transcript induction or repression to that observed during N-starvation for the majority of the DEGs which became even more similar during HL exposure (Fig. 6A). The transcriptional changes point to an altered N homeostasis induced by *ILP1*-deficiency (Fig. 6B). Root growth was strongly suppressed in WT plants grown in the absence of N in the media, compared to plants grown on commercially available MS medium or self-made MS medium supplemented with an N source (Fig. 6C). Retardation of root growth of WT plants under N-deficiency was as strong as for *ilp1-1* and *ntr1-1* mutants on regular MS (Fig. 6C). It is notable that the root length of *ilp1-1* was entirely insensitive to the N content of the media, but the *ntr1-1* mutant still showed a significant reduction of root length on -N media compared to the control plates. These results suggested that, as far as root growth is concerned, the spliceosomal complex mutants are in a status of nitrogen deficiency and that ILP1 could play a more dominant function than NTR1. However, high N content in MS media (40 mM total N) did not rescue the root growth of the *ilp1-1* and *ntr1-1* mutants (Fig. 6C), and when the mutants were grown on soil the regular application of fertiliser (0.2 g N l^-1^) did not suppress their pronounced anthocyanin accumulation (Fig. 6D). We also observed no significant differences in total C, N and protein content between WT and *ILP1*-deficient plants (Fig. 6F). The apparent and particular perturbation of AA biosynthesis in HL (Fig. 5) led us to examine the expression of genes associated with N assimilation and AA metabolism in more detail (Supplemental Fig. S8). Given that a pronounced accumulation of glutamate, the major amino group donor for AA biosynthesis, and diminished content for the majority of AA was observed in HL-treated *ilp1-1* and *ntr1-1*, we focused the analysis on aminotransferases and enzymes involved in NH_4_^+^ assimilation – glutamine synthetase (GS) and glutamine:2-oxoglutarate aminotransferase (GOGAT). From the targeted analysis of the RNAseq data, we found only minor differences in the expression of these genes. For instance, two enzymes involved in alanine biosynthesis were significantly induced (*ALAAT2*) or repressed (*GGAT2*) in *ilp1-1* compared to WT during the HL treatment (Supplemental Fig. S8A). Aminotransferases use pyridoxal 5’-phosphate (PLP, vitamin B6) as a co-factor for the transamination reaction. Among the gene products involved in PLP metabolism, two pyridoxal phosphate phosphatases, *PS2* and *PYRIDOXAL PHOSPHATE PHOSPHATASE-RELATED PROTEIN1* (*PEPC1*), displayed higher expression levels in *ilp1-1* compared to WT (Supplemental Fig. S8B), suggesting changes in PLP metabolism. We, therefore, considered the possibility of low PLP contents in the spliceosomal mutants being causative for the defect in AA biosynthesis. However, neither the defects in hypocotyl or root growth nor the overaccumulation of anthocyanins in HL-treated *ilp1-1* and *ntr1-1* were rescued by exogenous PLP application (Fig. 6F), indicating that PLP deficiency did not cause the metabolic restriction in the mutants.

**Figure 6:**
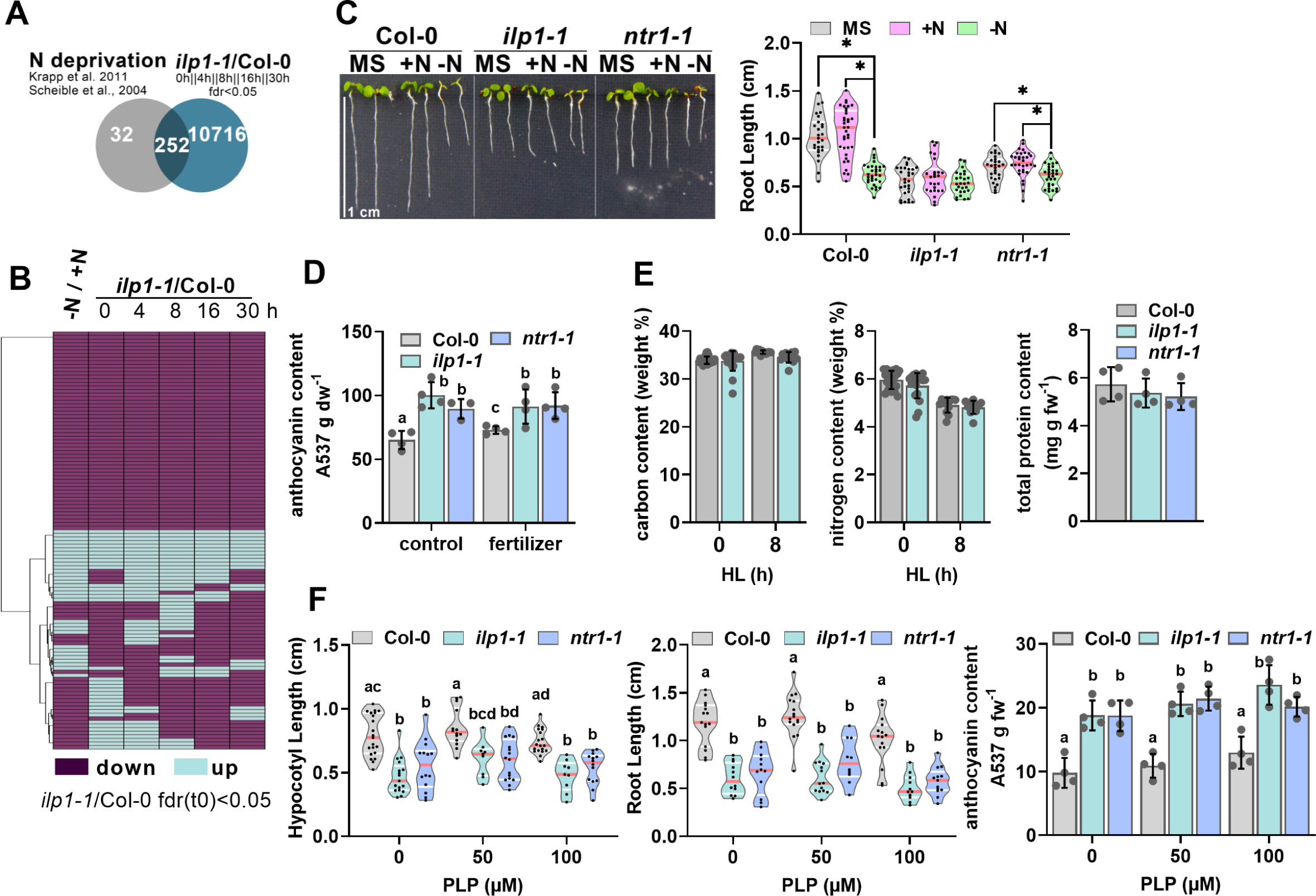
Mutants of *ILP1* and *NTR1* show symptoms of nitrogen (N) deficiency without being N-limited. (**A**) Overlap of differentially expressed genes in *ilp1-1* relative to Col-0 in at least one time-point of the HL kinetic (fdr<0.05) and transcripts de-regulated in response to N-deprivation (Krapp et al. 2011 and Scheible et al. 2004). (**B**) Heatmap showing relative expression changes of mRNAs in response to N-depletion (first column) and DEGs in *ilp1-1* relative to Col-0. Transcripts differentially expressed in *ilp1-1* at t0 (fdr(t0)<0.05) were included. Changes in expression relative to the control (+N or Col-0) were manually assigned +1 (blue, induced) and -1 (purple, repressed). (**C**) Root length of 7-DAG Col-0, *ilp1-1*, and *ntr1-1* (n≥27) in ½ MS, 100% nitrogen (40mM), and no nitrogen-containing media. Asterisks indicate statistical significance compared to Col-0 by Student’s *t*-test (\**p<*0.05). The red lines inside the violins show the median of values and the white lines the quartiles. (**D**) Anthocyanin contents of 4-week old Col-0, *ilp1-1*, and *ntr1-1* grown with or without fertilizer (0.2 N g l^-1^) after 24 h HL. Data are mean ± SD (n=4). The red lines inside the violins show the median of values and the white lines the quartiles. Different letters indicate significance groups at *p<*0.05 by Tukey’s multiple comparison test as determined by one-way ANOVA. Values with the same letter are not significantly different. (**E**) Total carbon and nitrogen contents in Col-0 and *ilp1-1* (n=12) at t0 and 8 h HL and total protein contents in 4-week old Col-0, *ilp1-1* and *ntr1-1*. (**F**) Hypocotyl length (4-DAG), root length (6-DAG) and anthocyanin content in 8-DAG seedlings grown in 12 h L/ 12 h D after 24 h HL exposure in the presence or absence of Pyridoxalphosphate (PLP). Different letters indicate significance groups at *p<*0.05 by Tukey’s and Šidák’s multiple comparison tests, respectively, as determined by two-way ANOVA.

## Discussion

Plant acclimation is underpinned by a multilayered adjustment of cellular processes to maintain an existing or establish a new cellular homeostasis permitting growth, reproduction or even survival in an ever-changing environment (Huang et al., 2019; Kleine et al., 2021; Schwenkert et al., 2022; Richter et al., 2023). In response to sudden fluctuations of environmental conditions, mRNA contents, followed by changes in the abundance of enzymes and post-translational regulation, control the biosynthesis and distribution of metabolites between different pathways to meet the demand for end products. By a forward genetic screen, we identified ILP1 and PRL1 as factors that influence anthocyanin accumulation in HL (Fig. 1, Supplemental Figs S1 and S3, and Supplemental Table S7), revealing that spliceosomal complex (components) are essential for the conditional and rapid adjustment of the transcriptome (Figs 3-4) and metabolome (Figs 5-6) when plants face a shift to high light intensities and, hence, are crucial factors for plant HL acclimation.

Our results showed that AA metabolism and other branches of primary and secondary plant metabolism depend on photosynthetic activity and the function of the spliceosomal complex components ILP1, NTR1, and PRL1 in HL. Mutants of these factors showed a conditional, i.e. only in HL, accumulation of Glu and low contents of Gln and other AA (Fig. 5, Supplemental Fig. S3, and Supplemental Table S8). Given that Glu is the primary amino group donor for AA biosynthesis (Forde and Lea, 2007), a perturbation of de-novo AA biosynthesis could be assumed. On the other hand, accelerated catabolism of AA or channelling towards other branches of metabolism depending on AA, such as glutathione biosynthesis (Noctor et al., 2012), in the spliceosome mutants is not excluded. However, at least WT-like photosynthesis (Supplemental Fig. S6) and high amounts of reduced carbohydrates (Fig. 5) excluded a starvation response necessitating AA breakdown for energy production. In the light, NH_3_ released from the photorespiratory pathway by GLYCINE (Gly) DECARBOXYLASE (GDC) during the conversion of Gly to Serine (Ser) is re-assimilated by cytosolic and plastidial GS (Keys et al., 1978). Hence, low Gln and high Glu levels in the spliceosome mutants could indicate a perturbation of either GS function or changes in photorespiration. A constitutive disturbance of GS activity and N-assimilation in *ilp1-1* and *ntr1-1* is excluded because N and protein contents and accumulation of abundant photosynthetic proteins was WT-like under standard growth conditions (Fig. 6 and Supplemental Fig. S6). Likewise, under normal light photorespiratory release of NH_3_ and re-assimilation did not limit AA biosynthesis in the spliceosome mutants (Fig. 5J and Supplemental Table S8). Photorespiration is vital for the detoxification of 2-phosphoglycolate, refilling of the Calvin–Benson cycle with carbon skeletons and serine biosynthesis in light-grown plants (Timm and Hagemann, 2020; Fu et al., 2023). A high Gly/Ser ratio usually indicates an increased demand for photorespiratory activity or a limitation in GDC activity, particularly in HL (Engel et al., 2007). While the Gly/Ser ratio increased by several folds upon HL exposure of WT, the increase was less pronounced in *ilp1-1*, *ntr1-1* and *prl1-2* (Supplemental Table S8). The low Gly/Ser ratio of the spliceosome mutants under HL could be explained by a reduced flux of carbon towards the photorespiratory cycle, and a low Gly level would support this assumption. However, increased activity of enzymes involved in photorespiration, such as the GDC, would explain low Gly and, even more importantly, the increased Ser contents in *ilp1-1* and *ntr1-1* after short-term HL exposure (Fig. 5J). It will be of further importance to reveal if the improved photosynthetic activity of *ilp1-1* under HL (Supplemental Fig. S6) is partially caused by an improved photorespiration.

A remarkable increase in cellular sugar contents was apparent in *ilp1-1* and *ntr1-1* under HL (Fig. 5), suggesting that the limitation of, e.g., N-dependent AA biosynthesis results in a backup and accumulation of carbon-skeletons and carbohydrates when photosynthetic activity and carbon-fixation is stimulated in HL (Fig. 5 and Supplemental Fig. S6). Pronounced sugar and starch accumulation is well documented for various species under constitutive N-limitation (reviewed in Hermans et al., 2006; Jezek et al., 2023). Our results revealed a close connection between N and carbon metabolism when the biosynthesis of N-containing compounds is restricted in HL. Notably, although the mutants showed symptoms of N-deficiency, such as a short root during early seedling growth or deregulation of N-responsive transcripts in adult plants (Fig. 6), photosynthetic activity, the abundance of photosynthetic proteins and content of N-containing chlorophylls (Supplemental Fig. S6) were WT-like in *ilp1-1* and *ntr1-1* under standard growth conditions. Furthermore, AA contents (Fig. 5) and total N or protein content (Fig. 6) were not affected by *ILP1*-or *NTR1*-deficiency in normal light. It is noteworthy that neither accumulation of carbohydrates nor perturbation of AA biosynthesis was observed in anthocyanin overaccumulating *pap1-D* or the *se1* mutant with defects in miRNA biogenesis (Fig. 5, Lobbes et al., 2006), ruling out an (in)direct effect of anthocyanin biosynthesis and (SE1-dependent) miRNA biogenesis on ILP1 and NTR1-dependent regulation of metabolism. The results exclude a constitutive alteration of N and carbon assimilation and metabolism in the mutants. Instead, the results reveal that spliceosomal complex function and components are vital for adjusting primary and secondary metabolism at the (post-) transcriptional and, most likely, the post-translational level, thereby maintaining the C:N balance during acclimation to HL. The finding that *ilp1-1* and *ntr1-1* show symptoms of N-deficiency without being N-depleted (Fig. 6) may point to a sensory function of these factors to activate AA biosynthesis when the carbohydrate content in the cell increases. However, further analysis is required to disentangle cause from consequence, i.e., to reveal if ILP1/NTR1-deficiency primarily affects sugar biosynthesis followed by a secondary effect on AA biosynthesis or vice versa. Also, the possibility of changes in PLP biosynthesis or integration into aminotransferases, directly or indirectly affected by the spliceosome function, should be considered in the future.

No direct connection between transcript abundance and splicing efficiency/defects at a global level was found across the analysed samples in WT and *ilp1-1* (Supplemental Fig. S5). At first glance, our transcriptome studies suggest that alternative splicing plays a minor role during HL acclimation in WT plants (Fig. 4A-C). However, in agreement with the known function of alternative splicing for cold acclimation (Calixto et al., 2018), we found that ILP1-dependent splicing exists in WT (Fig. 4F). Therefore, it is likely that splicing defects and expression changes of mRNAs for various metabolic pathways before and during HL exposure (Figs. 3-4) are causative for the conditional restriction of metabolic acclimation in the analysed spliceosomal complex mutants. Light-dependent alternative splicing involving phytochromes and cryptochromes (reviewed in Zhang et al., 2017) and plastid-derived signals depending on photosynthetic activity (Petrillo et al., 2014) have been found. Future studies employing targeted analysis of individual transcripts will reveal the impact of (ILP1-dependent) splicing on the plants’ HL response. However, in normal light conditions splicing defects in *ilp1-1* (Fig. 4) resulted only in minor metabolic alterations (Fig. 5), which supports the hypothesis that ILP1 function is needed for the conditional regulation of metabolism during acclimation.

Besides splicing, ILP1 also affects gene expression directly via a C-terminal GC-rich domain (Yoshizumi et al., 2006). Based on the time-resolved RNAseq analysis, it is suggested that the deregulation of several biosynthetic pathways (Fig. 3) is causative for the altered HL acclimation response in *ILP1*-deficient plants. Among others, sugar and flavonoid biosynthesis genes were upregulated before and during the HL shift. Hence ILP1 could function as a repressor of these pathways. However, *SUS2* (Fig. 3E) and anthocyanin biosynthesis genes were more pronounced induced in *ilp1-1* compared to WT, and expression of anthocyanin biosynthesis genes was restored to WT-like level in *ILP1* overexpression lines (Fig. 2C and Fig. S2). This suggests that ILP1 is not a direct transcriptional regulator (suppressor) of anthocyanin biosynthesis, but a suppressor function in sugar signalling, as previously proposed, is not excluded (Nemeth et al., 1998). Also, root and hypocotyl length of young seedlings were again WT-like in the ILP1 complementation lines with a different degree of transgene expression (Fig. 1). Hence, perturbation of growth in *ilp1* mutants is most likely not caused by ILP1 acting as a direct transcriptional activator of growth-related transcripts. Instead, the more pronounced and rapid activation of anthocyanin biosynthesis in *ilp1-1*, *ntr1-1* and *prl1-2* can be explained by the overaccumulation of sugars or sugar-phosphates, including signal and regulatory molecules such as Tre6P (Figs. 2, 3 and 5). We recently demonstrated the connection between photosynthesis, carbon fixation, cellular sugar contents and activation of anthocyanin biosynthesis in HL (Zirngibl et al., 2023). In the current model, the accumulation of Tre6P under HL suppresses SnRK1 activity, thereby permitting anthocyanin accumulation (Broucke et al., 2022; Zirngibl et al., 2023). Previously, Tre6P and other sugar-phosphates were identified as a repressor of SnRK1 activity and the upstream SnRK1 activating kinases (Zhang et al., 2009; Nunes et al., 2013; Zhai et al., 2018; Baena-Gonzalez and Lunn, 2020). We propose that the spliceosomal complex mutants showed a more pronounced activation of anthocyanin biosynthesis in HL because they accumulated sugars and Tre6P more rapidly than WT plants (Fig. 5), resulting in faster activation of anthocyanin biosynthesis genes, such as *PAP1* and *DFR* (Fig. 2D-F). This assumption is strongly supported by the fact that inhibition of photosynthetic activity suppressed carbohydrate and anthocyanin accumulation in both WT and, more importantly, *ilp1-1* (Fig. 5F-H). Based on the analysis of *se1,* we also assume that (SE1-dependent) miRNA production is insignificant for HL-induced activation of anthocyanin biosynthesis in leaves. For instance, miR156 targets a repressor of PAP1, the *SQUAMOSA PROMOTER BINDING PROTEIN-LIKE* (*SPL*) mRNA, thereby de-repressing anthocyanin biosynthesis genes (Gou et al., 2011). However, miR156 is markedly reduced in *se1* mutant (Lobbes et al., 2006) without affecting anthocyanin accumulation in HL (Supplemental Fig. S3D-E). This finding is another example illustrating that different mechanisms govern the regulation of plant primary and secondary metabolism depending on the tissue, developmental stage and growth condition and justifying the need for a detailed analysis of the plant acclimation response in the future.

Production of anthocyanins, and other flavonoids, is a highly carbon-demanding process, requiring carbon skeletons channelled through the shikimate and sugar biosynthesis pathways. One of the most abundant anthocyanins in Arabidopsis, A11, is glycosylated with three molecules of glucose and one xylose (Saito et al., 2013). Therefore, anthocyanin biosynthesis represents a substantial sink for carbon and carbohydrates in plant metabolism. Although direct experimental evidence is scarce, and this connection seems to not exist in cold-acclimated plants (Kitashova et al., 2023), it is a long-standing hypothesis that activation of anthocyanin biosynthesis in adverse growth conditions could serve as safety valve to sequester excessive carbohydrates when other carbon sinks are limited (e.g., AA or sucrose biosynthesis or export) (Hernandez and Van Breusegem, 2010; Lei et al., 2011; Wippel and Sauer, 2012; Jiang et al., 2020). In *ilp1-1* and *ntr1-1,* diminished consumption of carbon skeletons for AA biosynthesis or the restriction in N metabolism most likely resulted in carbohydrate accumulation in HL. A perturbation of sugar metabolism and overaccumulation of starch and sucrose after sucrose feeding is also reported for *PRL1*-deficient plants (Nemeth et al., 1998). It is tempting to assume that excessive carbon skeletons cannot be metabolised in *prl1-2* because of restriction in N-assimilation or metabolism (e.g., AA). It could be speculated that the sugar accumulation then induces anthocyanin biosynthesis to enable the flow of carbon building-blocks towards anthocyanin/flavonoid biosynthesis to balance the different metabolite pools. Indeed, though in a different context, Jiang et al. (2020) demonstrated that repression of metabolic/biosynthetic pathways results in the activation of anthocyanin biosynthesis genes and a re-direction of carbon towards flavonoids/anthocyanins.

In conclusion, we propose that ILP1, NTR1, and PRL1 are vital for the conditional regulation of gene expression and exert a positive function on the regulation of metabolism during HL-acclimation. Furthermore, the spliceosomal complex and its function are essential for the balanced distribution of metabolites in the plant cell when plants are challenged with HL.

## Methods

### Plant Growth Conditions

*Arabidopsis thaliana* plants (Columbia-0 ecotype) were grown under 10h light/14h dark regime at 22°C, unless stated otherwise. Murashige and Skoog medium (Duchefa, M0222.0050) was used at half-strength concentration for plant culture Murashige and Skoog, 1962. For root and hypocotyl length experiments, seeds were surface-sterilized with 70 % ethanol and then stratified at 4°C in the dark for 2−4 d before seeding. Plates were kept verically for root length experiments while plates were kept in the dark after 6 hours of light induction for the hypocotyl length experiments.

### Plant Materials

The following mutant seeds were obtained from NASC seed stock centre: *ilp1-1* (SALK_030650C), *ilp1-2* (SALK_135563C), *ntr1-1* (SALK_073187), *prl1-2* (SALK_008466C), *pap1-D* Borevitz, 2000 and *serrate1* (Ori, 2000;Prigge and Wagner, 2001). Homozygous plants were isolated by PCR-based genotyping with gene-specific primers and T-DNA-specific primers. For *se-1* mutants, DNAs were sent for sequencing to check for the 7bp deletion in the *SERRATE* genomic sequence. See Supplemental Table S9 for the oligonucleotide sequences used in this study.

### HL, cold, and DCMU Treatment

High light treatment of 4-week-old plants was previously described. Briefly, 2 h after the onset of light, plants were subjected to 500 μmol photons m^−2^ s^−1^ of continuous light for 24 h with constant temperature (22°C) in a Conviron GEN1000 (Canada) growth chamber equipped with a white LED light source. To block photosynthesis, a solution of 200 µM DCMU (3-(3,4-dichlorophenyl)-1,1-dimethylurea) in water was applied to the leaves of 4-week-old plants using a paint brush. For the cold experiment, 4-week-old plants were kept at 4°C for one week. Cold treatment was started five hours after the onset of light. Sampling was done on mid-day seven days after cold treatment.

### Harvesting of Samples

Experiments were done with up to four replicates containing material of three 4-to-5-week-old plants grown on individual pots. Samples were immediately frozen in liquid nitrogen and lyophilized before further processing. Dried, ground, and fine leaf material was used for downstream experiments unless otherwise stated. Each experiment was repeated at least once.

### Statistics

All data sets were tested for significant differences using appropriate statistical tests. Tests and results for each figure are provided in Supplemental Table S10.

### Plasmid Construction and Transgenic Plant Production

For *pILP1:ILP1 ilp1*, 6641 bp of the *ILP1* genomic region was amplified by PCR with the primers GA16 and GA17. The amplicon contained the 1499 bp upstream of the start codon (including the endogenous promoter) and 840 bp downstream of the stop codon. The fragment was cloned into the pCambia3301 vector using *Sma*I restriction sites. For the *p35S:ILP1-HA*, 2724 bp of the coding sequence (CDS) of ILP1 was amplified using primers GA18 and GA90, and was cloned into the *Sm*aI site of pCambiaStrep plasmid vector. The desired vectors were transformed into *ilp1-1* and Col-0 plants via *Agrobacterium* tumefaciens-mediated gene transformation (floral dipping). T1 transformants were selected by Phosphinotricine treatment and resistant plants were genotyped for the *ilp1-1* background (LP= GA13; RP= GA14; LB= GA01) and endogenous *ILP1* (GA38 and GA344). Presence of the transgene was confirmed using primers GA44 and GA26. Selection and genotyping were repeated in the T2 generation. The *ILP1* expression levels in the knockout mutant alleles, genomic complementation, and overexpression lines were checked using primers GA47 and GA48.

### Suppressor Screening

The suppressor screen was based on the *gun5-1* point mutant with a defect in Mg chelatase subunit H of the chlorophyll biosynthesis pathway. When germinated in the presence of Norflurazon, the *gun5-1* seedlings exhibited lower anthocyanin accumulation due to a defect in chloroplast-to-nucleus signaling activating nuclear expression of anthocyanin biosynthesis genes Richter et al., 2020. To isolate second-site suppressor mutants exhibiting a *restored anthocyanin accumulation* (*raa*) phenotype, *gun5-1* seeds were treated with Ethyl methanesulfonate (*EMS*). In the M2 generation after EMS treatment, seeds were surface sterilized and plated on half-strength MS plates containing 5 µM NF. After 5 d in continuous light (100 µmol photons m^-2^ and s^-1^) seedlings were visually screened for change in leaf coloration (anthocyanin accumulation indicated by a purple color of the cotyledons). Candidates were transferred to 0.5xMS plates without NF and supplemented with 1% sucrose (w/v) and recovered for 10-14 d in low light. After emergence of green leaves, plants were transferred to soil for seed production and further characterization. The selected *raa* mutants were backcrossed to the parent *gun5-1* and re-selected based on the phenotype in the T2 generation. Leaves of T2 plants were pooled and used for whole genome sequencing and SNP identification. The *raa20* mutant was also backcrossed to Col-0 to isolate the *ilp1^SNP^* mutant without *gun5-1* background.

### Anthocyanin Quantification

Anthocyanins were extracted from leaf material with 1 ml of anthocyanin extraction buffer (18 % 1-propanol and 1% hydrochloric acid in water). The mixture was thoroughly mixed and left to incubate in darkness at RT for 2 hours. Then, the samples were centrifuged for 15 minutes (maximum speed) at 4°C. The supernatants were transferred to cuvettes, and the absorption levels at 537, 650, and 720 nm were measured. To calculate the absorption of anthocyanins, the following formula was employed: (A537−A720) − 0.25×(A650−A720). Finally, the values were normalized based on either the fresh weight (fw) or dry weight (dw) of the samples.

### Genomic DNA (gDNA) Extraction and SNP identification

To isolate the gDNA for SNP identification, cetyltrimethylammonium bromide (CTAB) method was used. Fine leaf material from 4-week-old leaves was added with 300 µL CTAB Buffer (2 % CTAB, 1.4 M NaCl, 0.1 M Tris-HCl pH 8.0) and mixed thoroughly. Then, 300 µL of RNAse (10 mg/ml) was added to each sample and was incubated at 65°C for 30 minutes with occasional mixing. Samples were placed on ice to cool down and 300 µL of chloroform was added. Then, tubes were centrifuged at 4°C for 15 minutes (maximum speed). The aqueous layer was transferred to new tubes and the DNA was precipitated by adding 250 µL ice-cold 2-propanol. Subsequently, samples were incubated at -20°C for 30 minutes. After centrifugation at 4°C for 15 minutes (maximum speed), the supernatant was discarded and the pellet washed with 500 µl of 80 % ethanol followed by centrifugation at 4°C for 5 minutes (maximum speed). The ethanol was discarded and the pellet was air dried for 5 minutes. The gDNA was resuspend in 30 µl of ddH2O. The Integrative Genomics Viewer (IGV; Robinson et al., 2011) was used for visualization of sequencing reads.

Two μg of DNA was used to prepare 350-bp-insert libraries for 150-bp paired-end sequencing (Novogene Biotech, Beijing, China) on an Illumina HiSeq 2500 system (Illumina, San Diego, USA) with standard Illumina protocols. The sequencing depth was at least 7 G raw data per sample which corresponds to a more than 50-fold coverage of the Arabidopsis thaliana genome. After grooming FASTQ files, adaptors were removed with Trimmomatic (Bolger et al., 2014), reads were mapped with BWA (Li and Durbin, 2010) to the TAIR10 annotation with the parameters ’mem -t 4 -k 32 -M’, and duplicates were removed by SAMtools (Li et al., 2009) with the rmdup tool. Single nucleotide polymorphisms (SNPs) were identified using SAMtools (Li et al., 2009) with the following parameter: ’mpileup -m 2 -F 0.002 -d 1000’. Only those SNPs were kept that were supported by more than 4 reads and of which the mapping quality was >20. To identify the SNPs specific for *raa20*, the SNPs between *raa20* and *gun5-1* were compared. The resulting *raa20* -specific SNP list was subjected to the web application CandiSNP (Etherington et al., 2014) which generates SNP density plots. The output list of CandiSNP was screened for nonsynonymous amino acid changes and the G/C to A/T transitions that were likely to be caused by EMS, with a special focus on the chromosome with the highest SNP density with an allele frequency of 1. Potential candidates were analysed by PCR-base methods and sanger sequencing (see below).

### RNA Extraction and quantitative PCR

RNA was extracted from ground leaf material by adding 300 μl of cell lysis buffer (2 % SDS, 68 mM sodium citrate, 132 mM citric acid, and 1 mM EDTA). Then 100 μl of a DNA/protein precipitation solution (4 M NaCl, 16 mM sodium citrate, and 32 mM citric acid) was added. The samples were vortexed and kept on ice for 10 minutes, and subsequently centrifuged at 4°C for 10 minutes (maximum speed). Then 300 μl of the resulting supernatant was combined with 300 μl of 2-propanol to precipitate the RNA. The mixture was centrifuged at room temperature for 5 minutes (maximum speed) and the RNA pellets were washed with 800 μl of 75 % ethanol, centrifuged again, and air dried. The RNA was dissolved in 25 μl of RNAse-free H_2_O and stored at -80°C until further use.

For quantitative polymerase chain reaction (qPCR) analysis, 1.5 μg of DNase I-treated RNA was transcribed into complementary DNA (cDNA) using RevertAid reverse transcriptase (Thermo Fisher, USA) following the manufacturer’s instructions. The qPCR analysis was carried out using a CFX96-C1000 96-well plate thermocycler (Bio-Rad) with ChamQ Universal SYBR qPCR Master Mix (Absource Diagnostics, Germany) and 1 μl of diluted (1:5) cDNA was used as template. The relative gene expression was determined using the 2^-ΔΔC(t)^ method with *SAND* (AT2G28390) as reference gene. The primer sequences used for the qPCR analysis can be found in Supplemental Table S9.

For RNA sequencing (RNA-seq), the isolated RNA was treated with 2 µL (2 units) of DNase I and 3 µL of 10X DNAse buffer and was kept at 37° for 30 minutes. Then 70 µL of ddH_2_O, 50 µL of 7.5 M ammonium acetate, and 400 µL of 100% ethanol were added to the samples. The RNA was precipitated by centrifugation at 4°C for 20 minutes and washed with 800 µl of 70 % ethanol. RNA was resuspended in 20 µL ddH_2_O, stored at -80°C until further use or shipped on dry ice.

### RNA Sequencing

Three biological replicates for Col-0 and tpt-2 after 9 h and 18 h of HL treatment were analyzed. Messenger RNA was purified from total RNA using poly-T oligo-attached magnetic beads (PolyA-enrichment). After fragmentation, the first-strand cDNA was synthesized using random hexamer primers, followed by the second strand cDNA synthesis. The library was tested with Qubit and real-time PCR for quantification and bioanalyzer for size distribution detection. Quantified libraries were sequenced on Illumina platforms (NovaSeq, paired end, 150bp), according to effective library concentration and data amount (4GB raw reads). Raw data of fastq format were processed through in-house perl scripts for quality control. This step obtained clean data (clean reads) by removing reads containing adapter, poly-N, and low quality. At the same time, Q20, Q30 and GC content of the clean data were calculated. Paired-end clean reads were aligned to the reference genome using Hisat2 v2.0.5. Reference genome and gene model annotation files of the TAIR10 release were used (ensemblplants_arabidopsis_thaliana_tair10_gca_000001735_1). featureCounts v1.5.0-p3 was used to count the reads numbers mapped to each gene. FPKM (Fragments Per Kilobase of transcript sequence per Millions of base pairs sequenced) of each gene was calculated based on the length of the gene and reads count mapped to this gene. RNAseq-analysis was conducted with two biological replicates of Col-0 and *ilp1-1* consisting of in total six plants per timepoint. RNAseq sample information and quality analysis are provided in Supplemental Table S11.

Raw sequencing data and count tables were depositied on the NCBI Gene Expression Omnibus website (https://www.ncbi.nlm.nih.gov/geo) under the accession number GSE236872.

**For review process, please use the following credentials:**

https://www.ncbi.nlm.nih.gov/geo/query/acc.cgi?acc=GSE236872

**Reviewer access token: mlshcqegtfklbiz**

### Data normalization, statistical analysis, PCA

Time course count data were normed using the median of ratios method (Love et al., 2014). Statistical tests were performed using DeSeq2 with default parameters (Love et al., 2014). PCA was conducted using FSharp.Stats v0.4.5 (Venn et al., 2022) on normed transcripts that were nonzero in all measured samples.

### Thermodynamically motivated enrichment analysis (TMEA)

FPKM data of transcript replicates were averaged, logged to the base *e* and filtered for the presence of count data in at least 4/5 time points. No absence of counts was allowed in the first and last time point in Col-0 and *ilp1-1* time courses respectively. Transcripts, that passed the filtering and were present in Col-0 and *ilp1-1*, were appended to a single sequence of length 10. Functional annotations for MapMan and gene ontology terms were obtained from Usadel et al. (2009), Venn and Mühlhaus (2022), Ashburner et al. (2000) and GO-Consortium (2021). Thermodynamically motivated enrichment analysis was performed as described in Schneider et al., 2020 (using TMEA v0.6.0).

### Splicing analysis

Inclusion level and significances were determined using rMATS. For every transcript a two-way-anova (Venn et al., 2022) was applied on inclusion levels of both genotypes at all four splice variants (SE, RI, A3, and A5). For every transcript a Two-Way-ANOVA (Venn et al., 2022a) was applied on inclusion levels of both genotypes at all four splice variants (SE, RI, A3, and A5). The *p* values were corrected to false discovery rates (Benjamini Hocherg). An ontology enrichment was performed on all elements with fdr < 0.05 and fdr < 0.01 (Venn and Mühlhaus, 2022).

### Starch Extraction and Quantification

Starch was extracted from dried and finely ground leaf material using 80 % (v/v) ethanol supplemented with 1 μl of 20 mg/mL Ribitol, and was incubated at 80°C for 30 min. Subsequently, samples were centrifuged for 10 min at RT (maximum speed) and the supernatant was transferred to new tubes for soluble sugar quantification. The pellet was resuspended in 75 0μl of 0.5 M NaOH and was incubated at 95°C for 30 min. Then, 750 μl of 1 M CH_3_COOH was added to the solution. The starch was then digested by mixing 100 μl of the starch suspension with 100 μl of amyloglucosidase solution (1 mg/ml in 200 mM CH_3_COOH and 100 mM NaOH) for 2 h with shaking at 1000 rpm at 55°C. After that, 100 µl of the starch digestion in ddH_2_O was mixed with 200 μl of glucose oxidase reagent (1 mg glucose oxidase, 1.5 mg horseradish peroxidase [HRP], 5 mg dianisidine/HCl in 50 ml of 0.5 M Tris/HCl [pH 7.0] and 40% [v/v] glycerol). For the glucose standard curve, 0, 100, 250, 500, and 750 µM glucose standards were mixed with 200 μl of the glucose oxidase reagent. The standards along with the samples were incubated at 30°C for 30 min before stopping the reaction with 400 μl of 5 M HCl. Samples were centrifuged briefly before analyzing the absorbance of the samples at 540 and 720 nm using a 96-well plate reader (SpectraMax M2 microplate reader [Molecular Devices, USA]). The amount of starch was determined as glucose equivalents, and the glucose content was calculated using the standard curve.

For the visualization of starch contents *in situ*, chlorophyll pigments were first destained with 80 % (v/v) ethanol for at least 20 min at 80°C until the leaves became transparent or chlorophyll-free. The rosette leaves were subsequently incubated for 1 h with Lugol’s iodine solution and destained with H_2_O afterwards.

### Cytosolic sugar quantification

From the dried supernatant of the starch extraction, 65 µL of Pyridin/methoxylamine (20 mg Methoxylamine/ 1 ml Pyridine; Sigma) was added to the tubes. The samples were incubated at 30°C with shaking for 90 minutes and were centrifuged briefly. Then, 35 µl of N-Methyl-N-trimethylsilyl-trifluoroacetamide (MSTFA) was added to the samples, incubated at 65°C for 90 minutes, and centrifuged briefly. Glucose, fructose, and sucrose were quantified by Gas Chromatography/Mass Spectrometry (GC/MS) (Agilent Technologies, USA). Ribitol was used as internal standard and peak areas of the sugars were normalized to the dry weight of the samples. Data were evaluated using the Agilent GC ChemStation software package (USA).

### Sugar-Phosphate extraction and analysis

Tre6P and other phosphorylated intermediates were extracted with chloroform-methanol and quantified by high-performance anion-exchange chromatography coupled to tandem mass spectrometry, as described in Lunn et al., 2006 with modifications from Figueroa et al., 2016.

### ROS Staining

Contents of superoxide radical were visualized using nitro blue tetrazolium chloride (NBT, Sigma-Aldrich [USA], 93862). Single leaves or whole rosette leaves from 4-week-old plants were incubated in NBT staining solution (25 mM HEPES/KOH, 1 mg/ml NBT, pH 7.5). Then the leaves were vacuum infiltrated for 15–30 minutes and the leaves were subsequently incubated for 2 h in the dark at RT. Chlorophyll was removed with 80 % (v/v) ethanol for 20 min at 80°C in a water bath. Accumulation of hydrogen peroxide was visualized using 3,3′-diaminobenzidine (DAB, Merck-Millipore [USA], D8001). An hour prior to the staining, the DAB staining solution (20 mM Tris/acetate, 1 mg/ml DAB [Sigma-Aldrich], pH 5.0) was prepared. Then the leaves were vacuum infiltrated for 30 minutes and the leaves were kept for 24 h in the dark at RT. Chlorophyll was destained with 80 % (v/v) ethanol for 20 min at 80°C in a water bath.

### Metabolite Quantification

Metabolite quantification was carried out by liquid chromatography coupled to tandem mass spectrometry (LC-MS/MS) analysis using the LCMS-8050 system (Shimadzu, Japan). Approximately 2-3 mg of dried and ground leaf material were mixed with 100 µl of LC-MS buffer (150 µL chloroform, 350 µL methanol, and 1 µl of 1 mg/ml morpholinoethanesulfonic acid (MES, internal standard)) and 400 µl ice-cold LC-MS grade H_2_O were added to the samples. After mixing thoroughly samples were incubated at -20°C for 2 hours. Samples were centrifuged for 10 minutes at RT (maximum speed) and the aqueous phase was transferred to new tubes. The first pellet was extracted again with 400 µl of ice-cold LC-MS grade H_2_O, were vortexed, and centrifuged for 5 minutes (maximum speed). The aqueous phase was combined with the first supernatant and was dried overnight in Speed-Vac (Eppendorf Concentrator plus^TM^). The metabolites were quantified using multiple reaction monitoring (MRM) according to the values provided in the LC-MS/MS method and the LabSolutions software package (Shimadzu, Japan). Authentic standards for the amino acids (Merck, Germany) were used for calibration and peak areas were normalized to signals of the internal standard (MES). Data were evaluated using the Lab solution software package (Shimadzu, Japan). Values of metabolite quantification is made available in Supplemental Table S8.

### PAM Measurement

Four-week-old plants were dark-adapted for 20 minutes prior to PAM measurement (Imaging PAM, Walz, Effeltrich, Germany). Plants were exposed to a pulsed, blue measuring beam (1 Hz, intensity 4, gain 1, damping 1) to obtain the basal chlorophyll (Chl) fluorescence (F) in darkness and a saturating light flash (intensity 10) was applied to determine the maximum Chl (F) in the dark-adapted state (Fm) used for calculation of Fv/Fm. The PSII quantum yield [Y(II)=(Fm′−F)/Fm′], NPQ [NPQ=(Fm−Fm′)/Fm′], non-regulated energy dissipation Y(NO), and photochemical quenching (qP) were measured at 286 µmol photons m^−2^ s^−1^ of a blue LED light source.

### Light Response Curves

Fluorescence light response curves (0, 25, 50, 100, 250, 500, 750, 1000, 1500 µmol m^−2^ s^−1^) before (t0) and after HL (t24 h) were measured using a Li-Cor-6400 gas exchange system (LI-COR, Lincoln, NE, USA) using fully expanded leaved from 7-week-old plants. The following conditions were set: block temperature = 25°C; CO_2_ concentration = 400 ppm; flow rate = 300 µmol s^−1^; and relative humidity = 60 to 70%.

### Western Blot and total protein quantification

Total leaf proteins were extracted from finely ground leaf materials using a protein extraction buffer (PEB, 56 mM Na_2_CO_3_, 56 mM dithiothreitol, 2% w/v SDS, 12% w/v sucrose, and 2 mM EDTA). After resuspension with PEB, samples were incubated at 70°C for 20 min and centrifuged for 10 min at RT (maximum speed). Protein extracts were transferred to new tubes and were separated in an 8 % or 12 % polyacrylamide-SDS gels and blotted onto a nitrocellulose membrane. Membranes were blocked for 1 h with 4 % milk solution in TBS-T (50 mM Tris/HCl, 150 mM NaCl, pH 7.5; 0.1 % v/v Tween 20), and were incubated overnight with the primary antibodies in 1 % milk solution in TBS at 4°C. The next day, membranes were washed and incubated with the secondary antibody for 1.5 h at RT (goat anti-rabbit immunoglobulin G [IgG] coupled with HRP, 1:10 000 in 1 % milk solution, TBS). Signals were detected using Clarity Western ECL substrate (Bio-Rad, Germany) and an ECL Chemostar CCD camera (Intas, Germany). Antibodies were purchased from Agrisera (Sweden) and Bio-Rad (Germany).

### Chlorophyll Extraction

For chlorophyll extraction and quantification, 1 ml of 80 % acetone with 10 µM KOH in water was added to 1-5 mg of dry weight of samples. Samples were vortexed and incubated for 1 h at -20°C, then centrifuged for 15 min (maximum speed) at 4°C. The Supernatants were transferred to a new tube and absorbance was analyzed at 646, 663 and 720nm. Chlorophyll *a* and *b* were calculated using the formula:

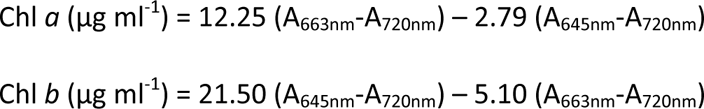

## Acknowledgement

Funding for the project was provided by the German Research Foundation (DFG) in the framework of the Collaborative Research Center/ Transregio “The green hub - Central Coordinator of Acclimation in Plants” (INST 86/2042-1/ project C06 to ASR, project C05 to DL, project C01 to TK, project D02 to TM). RF and JEL were financially supported by the Max Planck Society. We also thank Stefan Timm and Martin Hagemann (University of Rostock) for technical support with the LC/MS analysis and discussion, and Elena Heilmann and Peter Leinweber (Soil Science, University of Rostock) for C-and N-analyses and comments on the manuscript.

## Author contributions

ASR, conceptualization, supervision, project administration, experimentation. GEA, BV, NMK, RF, JL, TL, DL, TM, Investigation, data analysis, methodology, contribution to writing. ASR, TK, DL, JL, TM, acquisition of funding. ASR and GEA wrote the original draft with the contribution of all authors.

## Supplemental Figures

**Figure S1 (supports Figure 1): Suppressor screening of *gun5* for mutants with *restored anthocyanin accumulation* (*raa*) revealed *ILP1* (*raa20*) and *PRL1* (*raa7* and *14*) as factors determining anthocyanin accumulation.**

(**A**) Concept of the suppressor screen and phenotypes of *raa20*, 7 and 14 on Norflurazon-containing media (5d) and after 24 h high-light (HL) exposure.

(**B**) Melting peak analysis for the *ILP1* amplicon (primers from exon 4 and exon 5) in *gun5* and *raa20*. Note that compared to the *gun5-1* the different melting temperature for the *ILP1*-fragment amplified from *raa20* cDNA is indicative for a different length and composition.

(**C**) Relative expression of the retained intron in *raa20* using primers from intron 4 and exon 4. Col-0 samples served as control and *SAND* was used as reference gene. Data are mean ± SD (n=3).

(**D**) Schematic representation of the T-DNA insertions and position of the SNP in *ILP1* in *ilp1^SNP^*.

(**E**) *raa20* was backcrossed with Col-0 to isolate the *ilp1^SNP^* mutant.

(**F**) mRNA expression levels of the three *ilp1* mutants relative to Col-0 and *SAND* was used as reference gene. Data are mean ± SD (n=4).

**Figure S2 (supports Figure 2): *ILP1* mutant shows higher anthocyanin accumulation in cold and OE and genetic complementation lines display repression of transcripts 8 h after HL compared to *ilp1-1*.**

(**A** and **B**) Phenotypes (A) and anthocyanin accumulation (B) of 4-week old (SD conditions) Col-0, *ilp1-1*, OE and complementation lines after 7 days of cold treatment at 4°C. The start of cold treatment was set 5 h after the onset of light.

(**C** and **D**) Relative expressions of *PAP1* (C) and *DFR* (D) at 8 h after HL in 4-4-week old Col-0, *ilp1-1*, OE and complementation lines. Transcripts were compared relative to Col-0 at t0 and *SAND* was used as reference gene (n=4). Data are mean ± SD. Different letters indicate significant difference at *p<*0.05 by Tukey’s multiple comparison test as determined by one-way ANOVA. Values with the same letter are not significantly different.

**Figure S3 (supports Figure 2). Mutant of *PRL1*, another component of the spliceosome, resembled *ilp1-1* and *ntr1-1* during HL acclimation.**

(**A**) Hypocotyl lengths of 4-day old dark grown Col-0, *ilp1-1*, *ntr-1,* and *prl1-2* (n≥29). Asterisks indicate statistical significance compared to Col-0 by Student’s *t*-test (\**p<*0.05). The red lines inside the violins show the median of values and the white lines the quartiles.

(**B**) Root lengths of 7-day old Col-0, *ilp1-1*, *ntr1-1,* and *prl1-2* (n≥31) under SD conditions, respectively. Asterisks indicate statistical significance compared to Col-0 by Student’s *t*-test (\**p<*0.05). The red lines inside the violins show the median of values and the white lines the quartiles.

(**C**) Anthocyanin contents of 4-week old Col-0, *ilp1-1*, and *prl1-2* after 24 h of HL exposure. Data are mean ± SD (n=4). Asterisks indicate statistical significance compared to Col-0 by Student’s *t*-test (\**p<*0.05).

(**D**) Fructose (Fru), glucose (Glu) and sucrose (Suc) contents of *ilp1-1* and *prl1-2* before and 8 h after HL. Data are mean ± SD (n=4). Asterisks indicate statistical significance compared to Col-0 at each time point by Student’s *t*-test (\**p<*0.05).

(**E**) Log_2_ fold changes of the different amino acids compared to Col-0 before and after 8 h HL exposure. Data are mean ± SD (n=4). Asterisks indicate statistical significance compared to Col-0 by Student’s *t*-test (\**p<*0.05).

(**F**) (∑Amino Acids – Glu)/Glu in *ilp1-1* and *prl1-2* before and 8 h after HL. Data are mean ± SD (n=4). Different letters indicate significance groups at *p<*0.05 by Tukey’s multiple comparison test as determined by two-way ANOVA.

**Figure S4 (supports Figure 3): Gene ontology (GO) term enrichment of transcripts belonging to the constraints (C) 1-3 of the TMEA analysis.**

(**A** and **B**) C1 (pos and negative).

(**C** and **D**) C2 (pos and negative).

(**E** and **F**) C3 (pos and negative). GO term enrichment analysis was obtained from TMEA analysis with transcripts contributing to C1-3 in both genotypes. Corrected *P*-values from BH-analysis of 0 are shown as -log_10_(p-value) = 3. (G) Venn diagram and GO-term enrichment of DEGs in *ilp1-1*/Col-0 (fdr<0.05) at t0 (0h), 4 h and 8 h HL exposure.

**Figure S5 (supports Figure 4):** (**A**) Course of differentially splicing in Col-0 and *ilp1-1* across the HL kinetic. The number of transcripts (count) with corresponding inclusion level (IncLvl) of Retained Intron (RI), Skipped Exon (SE), and Alternative 3’ and 5’ ends (A3+A5) are plotted.2

(**B**) Correlative analysis of expression level (log(fpkm+1)) and IncLevel of differentially spliced transcripts.

**Figure S6 (supports Figure 5): *ILP1-*deficient plants show increased photosynthesis in HL.**

(**A**) Starch contents (expressed as glucose equivalents) of 7-week old Col-0 and *ilp1-1* under a normal day/night cycle (SD). Data are mean ± SD (n=4). Asterisks indicate statistical significance compared to Col-0 by Student’s *t*-test (\**p<*0.05). EoD, End of Day. EoN, End of Night

(**B**) Rates of starch degradation and starch synthesis in 7-week old Col-0 and *ilp1-1* under a normal day/night cycle (SD). EoD, End of Day. EoN, End of Night

(**C**) Left panel: Fv/Fm, coefficients of photochemical quenching (qP), non-photochemical quenching (NPQ), non-regulated energy dissipation Y (NO), and PSII quantum yield (Y(II)) of Col-0 and *ilp1-1* at t0, 8 h and 24 h after HL using an Imaging-PAM system (Walz) and are displayed on a rainbow color scale. Right panel: Quantitative analyses from six plants per genotype and six leaves (n = 36) were analyzed at 286 µmol photons m^−2^ s^−1^. Data are mean ± SD. Asterisks indicate statistical significance compared to Col-0 by Student’s *t*-test (\**p<*0.05).

(**D**) Net photosynthetic CO_2_ uptake rates of 7-week old Col-0 and *ilp1-1* (n≥12) in response to increasing light intensities (0, 25, 50, 100, 250, 500, 750, 1000, and 1500 µmol photons m^−2^ s^−1^) at t0 and 24 h after HL. Data are mean ± SD. Asterisks indicate statistical significance compared to Col-0 by Student’s *t*-test (\**p<*0.05).

(**E**) Total chlorophyll contents of 4-week old Col-0 and *ilp1-1*. Data are mean ± SD (n=4).

(**F**) Steady-state levels of photosynthetic proteins in Col-0, *ilp1-1*, and *ntr1-1* at t0, 8 h, and 24 h HL.

**Figure S7 (supports Figure 6): *ILP1* knockout mutants have WT-like ROS response and transcriptional landscape of phytohormones during HL.**

(**A**) Kinetic analysis of cytosolic *APX1* during the course of HL acclimation and de-acclimation in 4-week old Col-0 and *ilp1-1* mutants. Transcripts were compared relative to Col-0 at t0 and *SAND* was used as reference gene (n=4). Data are mean ± SD. Asterisks indicate statistical significance compared to Col-0 by Student’s *t*-test (\**p<*0.05). For the expression analysis of transcripts shown as FPKM, values were extracted from the RNAseq analysis (n=2 samples per time-point).

(**B** and **C**) H_2_O_2_ and superoxide (O_2_^●-^) staining by DAB and NBT, respectively, in 4-week old Col-0 and *ilp1-1* mutants during HL.

(**D**) Expression profiles of genes that are involved in hormone biosynthetic pathways under HL. The red lines inside the violins show the median of values and the white lines the quartiles. The y-axis is Log_2_ fold change (Log_2_ FC *ilp1*/Col-0). ABA (n=50), Auxin (n=36), cytokinin (n=29), GA (n=17), ethylene (n=22), JA (n=23), BR (n=10), and SA (n=7).

**Figure S8 (supports Figure 6):** Expression analysis of (**A**) aminotransferases and gene products involved in N-assimilation and (**B**) enzymes involved in PLP biosynthesis. For (B), enzymes listed in the KEGG database were analysed (https://www.kegg.jp/pathway/ath00750).

